# β-cell dedifferentiation is associated with epithelial-mesenchymal transition triggered by miR-7-mediated repression of mSwi/Snf complex

**DOI:** 10.1101/789461

**Authors:** Tracy CS Mak, Yorrick von Ohlen, Yi Fang Wang, Eva Kane, Kaste Jurgaityte, Pedro Ervilha, Pauline Chabosseau, Walter Distaso, Victoria Salem, Alejandra Tomas, Markus Stoffel, Piero Marchetti, A.M. James Shapiro, Guy A. Rutter, Mathieu Latreille

## Abstract

β-cell dedifferentiation has been revealed as a pathological mechanism underlying pancreatic dysfunction in diabetes. However, little is known on the genetic and epigenetic changes linked with the dedifferentiation of β-cells. We now report that β-cell dedifferentiation is associated with epithelial to mesenchymal transition (EMT) triggered by miR-7-mediated repression of Smarca4/Brg1 expression, a catalytic subunit of the mSwi/Snf chromatin remodeling complexes essential for β-cell transcription factors (β-TFs) activity. miR-7-mediated repression of Brg1 expression in diabetes causes an overall compaction of chromatin structure preventing β-TFs from accessing and transactivating genes maintaining the functional and epithelial identity of β-cells. Concomitantly, loss of β-cell identity impairs the ability of β-TFs Pdx1, Nkx6-1, Neurod1 to repress non-β-cell genes enriched selectively in mesenchymal cells leading to EMT, change in islet microenvironment, and fibrosis. Remarkably, anti-EMT agents normalized glucose tolerance of diabetic mice, thus revealing mesenchymal reprogramming of β-cells as a novel therapeutic target in diabetes. This study sheds light on the genetic signature of dedifferentiated β-cells and highlights how loss of mSwi/Snf activity in diabetes initiating a step-wise remodeling of epigenetic landscapes of β-cells leading to the induction of an EMT process reminiscent of a response to tissue injury.

## Introduction

Pancreatic β-cells are specialized epithelial cells within islets of Langerhans that secrete insulin to promote peripheral glucose uptake and meet metabolic demands (1). Deficiency of functional insulin-producing β-cell mass leads to a rise in glycemia and underlies the rising prevalence of diabetes worldwide (WHO 2019). Type 1 diabetes (T1D) is associated with auto-immune destruction of β-cells, whereas type 2 diabetes (T2D) is associated with gradual deterioration of pancreatic β-cell function and survival induced by obesity and insulin resistance (2, 3). The impairment in glucose-stimulated insulin secretion has been attributed to changes in glucose sensing (4) and mitochondrial dysfunction (5) whereas prolonged endoplasmic reticulum and oxidative stress triggers β-cell apoptosis (6). Moreover, genome-wide association studies (GWAS) have revealed that genetic alteration in multiple genes encoding crucial transcription factors regulating pancreatic β-cell differentiation (e.g. Sox7, Glis3, Hnf4*α*, Tcf7l2), glucose sensing (e.g SLC2A2, GCK, KCNJ11, ABCC8) or insulin packaging and secretion (SLC30A8, CDKN2A/B) may predispose individuals to pancreatic β-cell dysfunction in T2D (7–10).

Recent studies have however challenged our view of the prevalent pathogenic drivers of β-cell dysfunction in T2D. Indeed, β-cell dedifferentiation has been revealed as an alternative mechanism to β-cell apoptosis in T2D (11). Dedifferentiated β-cells loose insulin expression as well as other markers of functional identity including transcription factors and regulators of glucose-stimulated insulin secretion. Loss-of-function studies in mice have revealed that inactivation of a subset of β-cell transcription factor (β-TFs) (e.g. Pdx1, Nkx6-1, Nkx2.2, Pax6, Rfx6) (12–16) and epigenetic regulators (e.g. DNMT1, LIMD1 and EED/PRC2)(17–19) driving β-cell lineage development during embryogenesis is sufficient to trigger the dedifferentiation of mature β-cells and their transdifferentiation into other endocrine cell types (11, 12, 15, 20–24). High metabolic demand during pregnancy (11), insulin resistance (25, 26), inflammation (11, 27) and viral infection (28) have been shown to trigger β-cell dedifferentiation. Deciphering the genetic and epigenetic signature of dedifferentiated β-cells, thus represent a fundamental step toward the development of therapeutic agents fostering β-cell identity in T2D.

MicroRNAs (miRNAs) are crucial post-transcriptional regulators of gene expression which have been shown to play essential roles in cell lineage determination in many organs by modulating transcription factor expression and signaling networks that reinforce cellular decisions (29). Dysregulation of miRNA expression has been shown mediate stress responses in several disease including metabolic disorders (30, 31), suggesting that miRNAs could serve as targets for therapeutic intervention. We previously reported increased miR-7 levels in islets of rodent models of diabetes (e.g. *db/db* mice and non-obese GK rats)(32, 33) as well as in human islets transplanted into mice exposed to an obesogenic diet (32). The pathological significance of these observations was demonstrated following the generation of a β-cell-specific miR-7 overexpressing mouse model (Tg7) which develops early-onset diabetes due to β-cell dedifferentiation (32). Conversely, how elevated miR-7 levels in diabetes impair β-cell identity remains unclear.

By profiling gene expression and open chromatin regions of Tg7 islets, we now report that loss of β-cell identity triggers an islet injury response characterized by an epithelial to mesenchymal transition (EMT) process in both mouse and humans. Elevated miR-7 levels trigger the dedifferentiation of mature β-cells through the repression of Brg1/Smarca4, a catalytic subunit of mSwi/Snf chromatin remodelling complexes required for the transactivation lineage-specific and epithelial genes by β-cell transcription factors (β-TF) function. Our findings highlight how dysregulation miRNA networks can compromise the functional and epithelial identity of β-cells and trigger an EMT-like reprogramming process with therapeutic relevance in diabetes.

## RESULTS

### Loss of β-cells identity impairs islet connectivity in Tg7 mice

Genetic overexpression of miR-7 in β-cells results in diabetes from 4 weeks of age and is associated with decreased expression of mature β-cell differentiation markers, most notably *INS1/2* and β-TFs genes (Supplementary Figure 1). How dedifferentiation affects Ca^2+^ dynamics and long-range β-cell:β-cell connectivity in diabetes remains unclear. To this end, we monitored intracellular free Ca^2+^ dynamics in intact islets in response to elevated concentrations of glucose using a trappable intracellular Ca^2+^ dye and multicellular imaging. Ca^2+^ dynamics in response to 17 mM glucose were almost completely eliminated in Tg7 islets, whilst those in response to depolarisation with KCl were less affected (Figure1a-d). Moreover, assessment of cellular coordination using Pearson correlation analysis in Tg7 islets (34, 35) revealed that the glucose-induced increment in β-cell connectivity was sharply reduced in Tg7 *versus* Wt islets (Figure 1e-f). Supporting these results, we found that Tg7 islets displayed significantly decreased expression of Cx36 (Gjd2), a component of gap junction in epithelial cells, implicating this change as a possible contributor to the abrogated cell-cell connectivity observed in Tg7 islets (Figure 1g). In contrast, expression of voltage-gated calcium channel (VGCC) isoforms CaV1.2, CaV1.3, CaV2.1, CaV2.2, CaV2.3, and CaV3.1 was largely unchanged in mutant mice (data not shown). Whilst not directly interrogated at the imaging speed and with the analytical approaches deployed here (Pearson R correlation – (Hodson et al. 2013)), these data may be compatible with a loss of “hub-like” β-cells that orchestrate calcium oscillation islet-wide in wild-type islets (36). In any case, these results indicate that miR-7-mediated β-cell dedifferentiation is associated with both glucose-induced Ca^2+^ dynamics and long-range β-cells connectivity.

**Figure 1.**
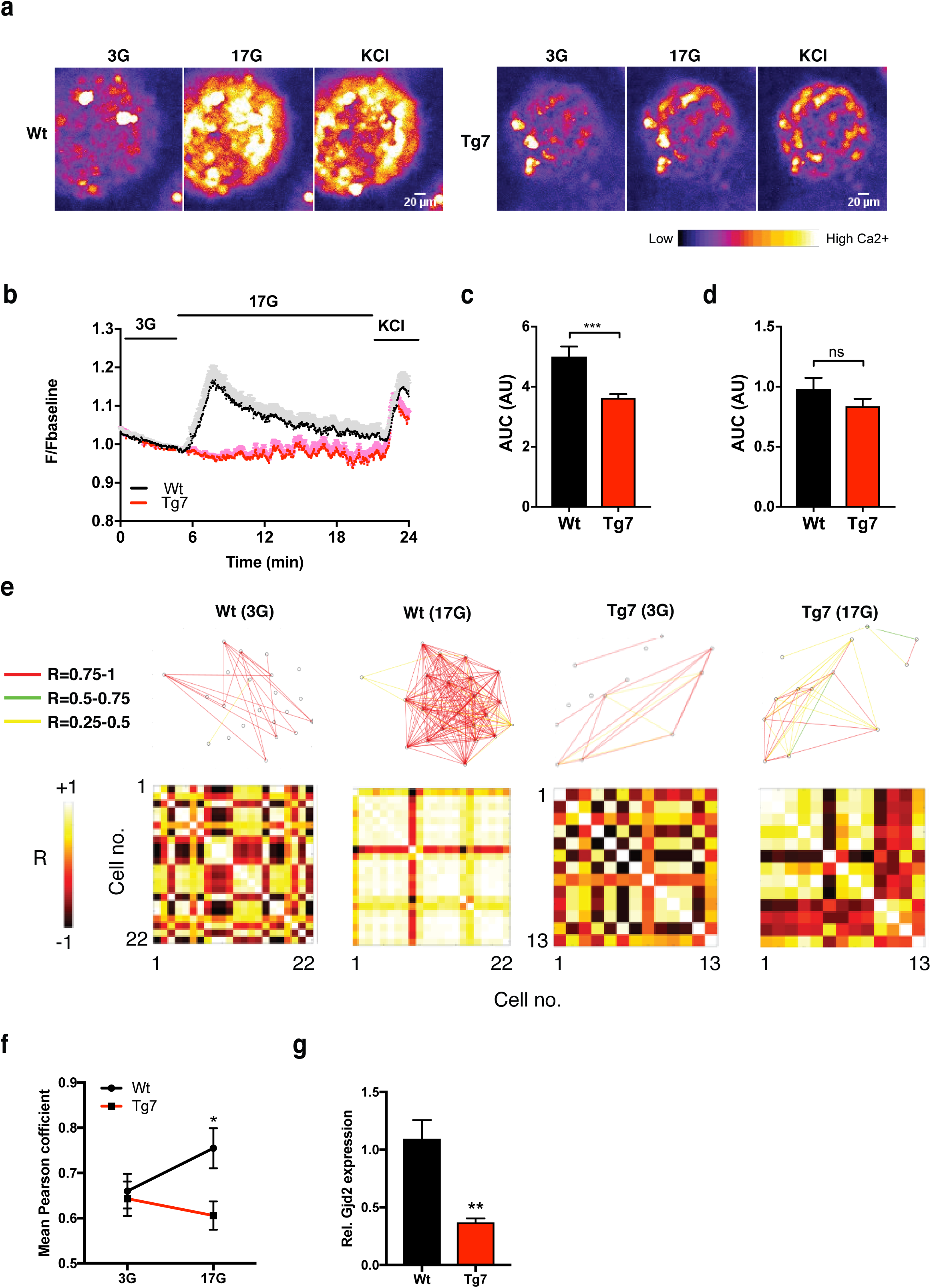
Dedifferentiated β-cells display impaired connectivity. (A) Representative images of pseudocolored 12-(12w) week-old Tg7 islet after exposure to permissive glucose concentrations (G3: 3 mM glucose; G11: 11 mM glucose; KCl: 20 mM); recording time: 40–60 minutes (B) Mean fluorescence intensity of Ca^2+^ oscillation in 12w Wt and Tg7 islets following elevation of glucose concentration and exposure to KCl. Individual traces were normalised (F/F baseline) and then averaged (C-D) Area under the curve (AUC) of each trace of cell from islets exposed to glucose (C) or KCl (D). n = 3 mice per genotype, from 2 independent experiments. Number of islets used: Wt: 23 and Tg7: 24. (E) Cartesian functional connectivity maps in 12w Tg7 islets displaying the x-y position of analysed cells within the imagedislet (black dots). Cells are connected with a coloured line if the p statistic for the Pearson coefficient wasp< 0.001 post bootstrapping. The strength of the cell pair correlation (the Pearson R statistic) was colour coded: red for R of 0.75 to 1.0, yellow for R of 0.5 to 0.75 and green for R of 0.25-0.5. Bottom: Heat maps show the Pearson coefficient of each cell pair in a colour-coded manner - negative correlation; dark brown (−1), no correlation; mid brown (0), high correlation; yellow/white (1). (F) Pooled data for the Wt (n=23) andTg7 (n=24) islets imaged in the connectivity analysis. (G) Expression of Gjd2/Connexin36 in 12w Tg7 islets.(n=7-8) Unpaired Student’s t-test unless stated otherwise. Data are means ± SEM, *p < 0.05, ***p < 0.001

### Transdifferentiation of β-cells into δ-cells in Tg7 mice

Given that loss of β-cell identity has been associated with the transdifferentiation of β-cells into other endocrine cell types (11, 12, 15, 20, 21), we quantified the incidence of Ins^+^ cells that express non-β-cell markers such as glucagon (Gcg) and/or somatostatin (Sst)(so-called polyhormonal cells) in Tg7 mice. Our analysis revealed that 2w Tg7 mice displayed an increased number of Ins^+^Gcg^+^ (1.1 % ± 0.3 in Wt vs vs 3.8 % ± 1.1in Tg7, p=0.041, n=5 per group) and Ins^+^Sst^+^ (2.4 % ±0.5 in Wt vs 4.3 % ± 0.6 in Tg7, p=0.046) polyhormonal cells despite maintaining normoglycemia (Figure 2a). Interestingly, and unlike Ins^+^Gcg^+^ polyhormonal cells, the number of Ins^+^Sst^+^ cells was significantly increased in 12w diabetic Tg7 mice compared to age-matched controls (1.7 % in wt vs 3.9 % in Tg7, p=0.01, n=5 per group) and this translated into increased δ-cell mass in diabetic mice (7.3%±0.6 in Wt vs 10.1%±1.1 in Tg7, p=0.02)(Figure 2b). These observations strongly suggest that elevated miR-7 levels promote the conversion of β-cells to δ-cells.

**Figure 2.**
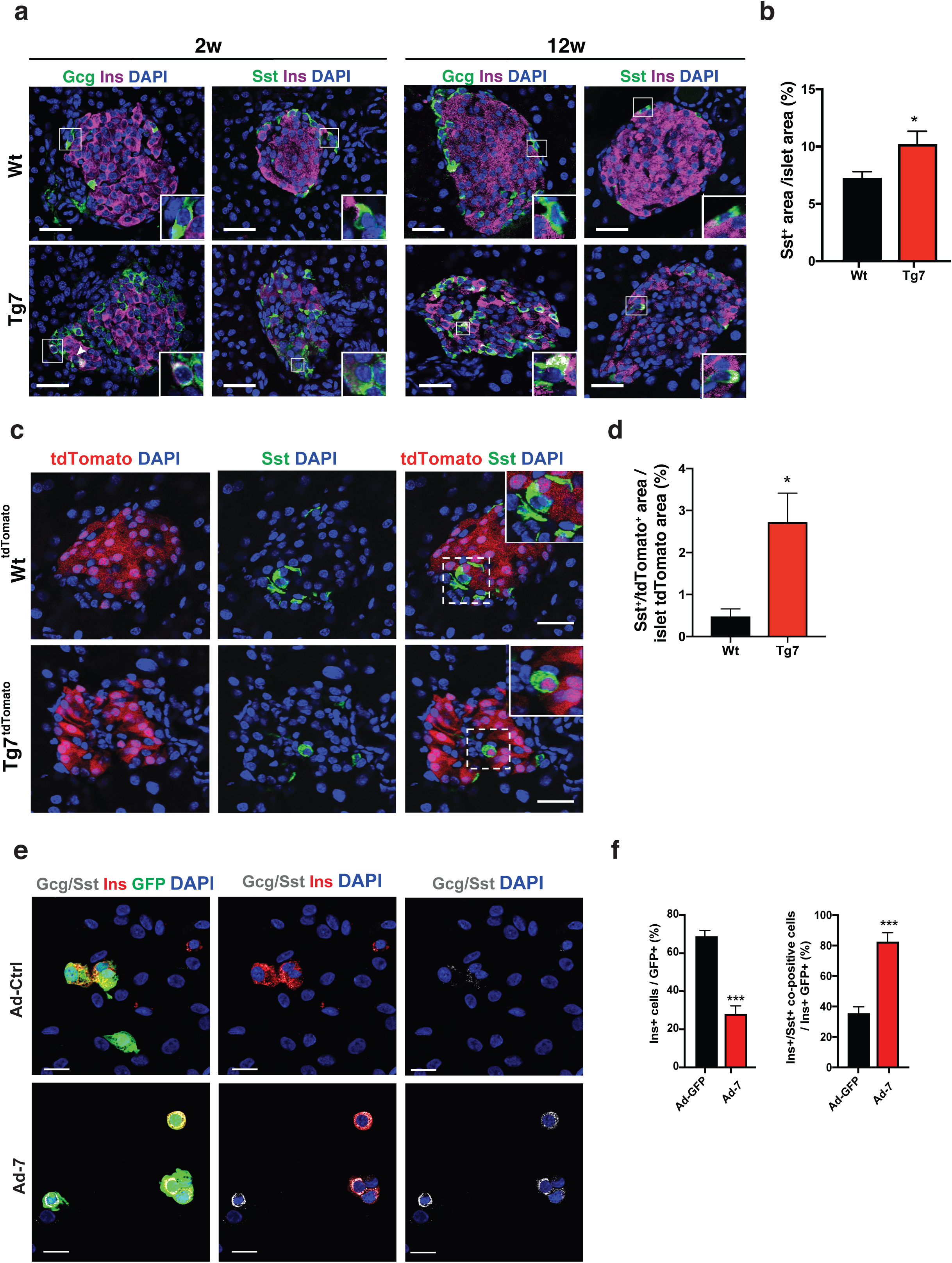
miR-7 overexpression triggers the transdifferentiation of β-cells into δ-cells inmice and humans. (A) Insulin and glucagon (Gcg) or somatostatin (Sst) co-immunofluorescence in pancreatic sections from 2-(2W) and 12-(12w) week-old Tg7 mice and Wt controls. Scale bar: 30mm (B) δ-cell mass calculated as the Sst^+^ area over DAPI^+^ islet area in Wt and Tg7 pancreas (n=5/group) (C) Genetic lineage tracing in tdTomato-labelled β-cells of 12w Wt and Tg7 mice. Shown is somatostatin (Sst) immunofluorescence pancreatic sections from Wt^tdTomato^ and Tg7^tdTomato^ mice. (D) Quantification of Sst^+^tdTomato^+^ cells in islets from D (n=5/group). (E) Insulin (red) and Gcg/Sst (white) cocktail co-immunofluorescence of dissociated human islets infected with Ad-Ctrl or Ad-7 adenoviruses. Infected cells were revealed by EGFP fluorescence (green) and nuclei by DAPI (blue) staining. Scale bar: 20mm. (F) Quantification of insulin^+^ and Ins^+^/Sst^+^ in EGFP^+^ cells shown in E. Data are means ± SEM, Unpaired Student’s t-test unless stated otherwise *p < 0.05, ***p < 0.001

To directly test this, we performed lineage tracing experiments in Tg7 mice carrying a Rosa26 tdTomato^Ripcre^ reporter and followed the fate of tdTomato-labeled β-cells. We found that the tdTomato^+^ area was similar in both genotypes (data not shown). Tracing analysis revealed that a significantly larger number of tdTomato-labelled β-cells from diabetic Tg7^tdTomato+^ mice expressed Sst (Figure 2c-d), thus demonstrating that miR-7 overexpression in β-cells promotes their conversion into δ-cells. Supporting this view, we found that infection of dissociated human islets with Ad-miR-7 increased the number of Ins^+^/Sst^+^ polyhormonal cells (Figure 2e-f). Indeed, more than 80 % of human Ins^+^ β-cells coexpressed Sst following miR-7 overexpression compared to 38 % in control infected islet preparations. These results indicate that induction of miR-7 triggers a β-to δ-cells transdifferentiation in both mouse and humans.

### Genetic signature of dedifferentiated β-cells from Tg7 mice

As a first step toward determining the molecular mechanism underlying dedifferentiation and transdifferentiation of β-cells in diabetes, we performed genome-wide RNA sequencing (RNA-seq) and assay for transposase-accessible chromatin using sequencing (ATAC-seq) analyses on islets isolated from non-diabetic 2w and diabetic 12w Tg7 mice (Figure 3a-b and Supplementary Figure 2a). A total of 2471 genes were differentially regulated in 2w islets (211 genes downregulated and 735 upregulated with at least a 2-fold change (2-FC), padj<0.05), whereas this reached 4224 in 12w Tg7 islets (348 downregulated and 1110 upregulated by at least 2-FC, padj<0.05)(Figure 3c, Supplementary Figure 2b). Gene set enrichment analysis (GSEA)(37) revealed that genes downregulated in 2w Tg7 islets display an overrepresentation of core β-cells components, including the *Unfolded Protein Response* (UPR) (*ERO1B*, *ATF6*, *XBP1*, *DNAJC3 WFS1*, *EDEM1*), *Pancreatic β-cell* identity (*UNC3*, *INS1*, *INS2*, *NKX6.1*, *MAFA*, *NEUROD1*, *INSM1*, *G6PC2*, *SLC2A2*, *ABCC8*, *SLC30A8*, *SYT4*), and *Protein Secretion* (*RAB2A*, *SNAP23*, *VAMP3*) (Figure 3d and Supplementary Table 1). In keeping with the ability of miR-7 to trigger β-cell dedifferentiation, we found increased levels of several β-cell “disallowed” genes normally repressed in mature β-cells (Figure 3e-f)(38, 39) and reawakening of pancreatic endoderm genes (Figure 3g-h). Of note, the expression of a significant proportion of differentially expressed genes found in 12w hyperglycemic Tg7 islets was already altered in 2w old non-diabeticTg7 islets. These results indicate that β-cell dedifferentiation represents a key pathological mechanism driving the early stages of pancreatic dysfunction in diabetes, challenging previous studies suggesting that β-cell dedifferentiation and reprogramming occurs secondarily to hyperglycemia (11, 21).

**Figure 3.**
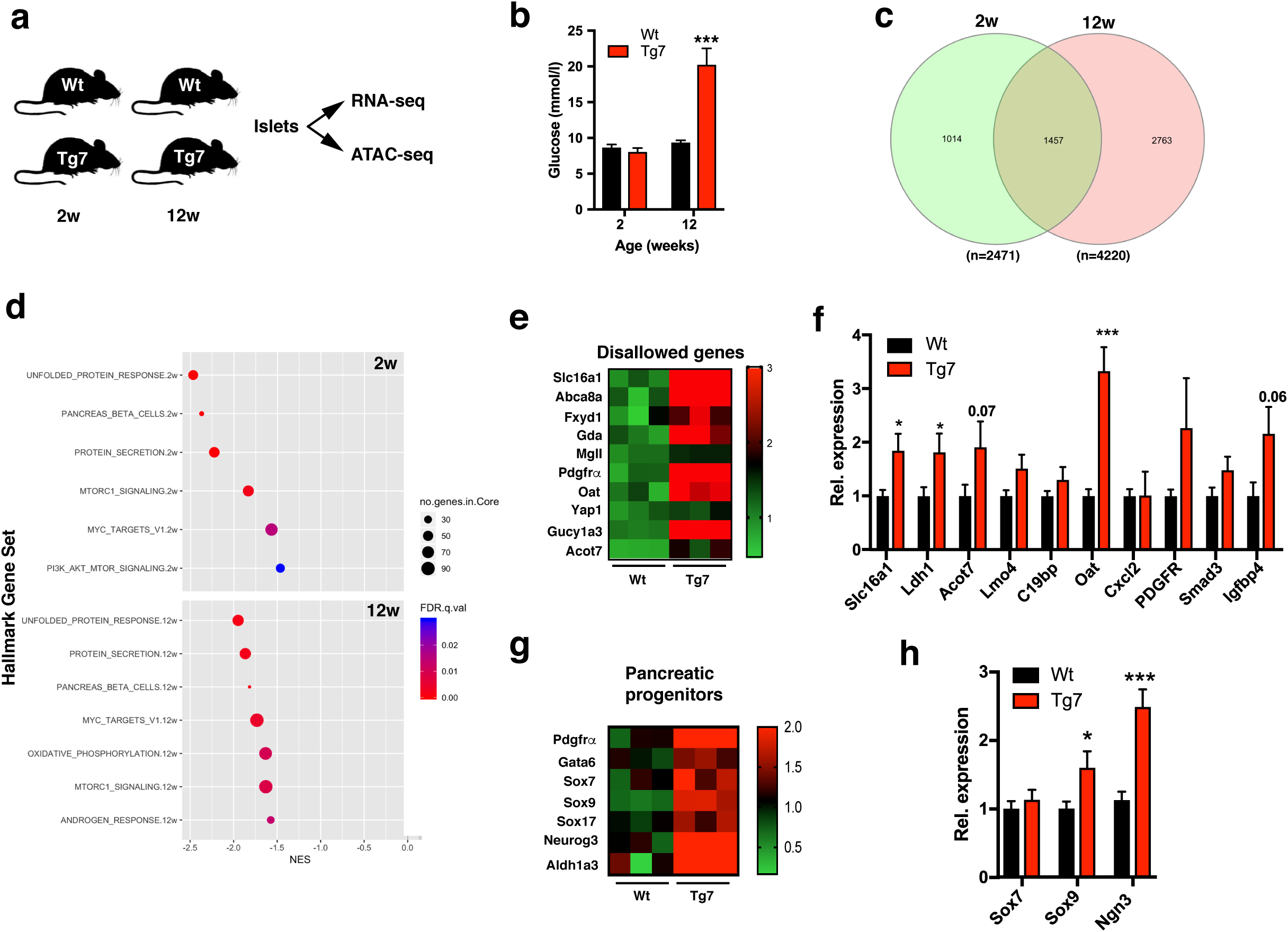
Genome-wide mRNA profiling of Tg7 islets. (A) Experimental overview. Islets were isolated from 2-(2w) and 12-week-old (12w) Wt and Tg7 mice and RNA subjected to RNA-seq and ATAC-seq (B) Random fed glycemia of Wt and Tg7 mice used in the RNA-seq (C) Venn diagram of differentially expressed genes in islets of 2w and 12w Tg7 mice p<0.05 (D) Gene set enrichment analysis (GSEA) of MSigDB Hallmark genes in downregulated pre-ranked gene ratios (Tg:Wt) from 2w (top) and 12w (bottom) Tg7 islets. Gene sets with indicated FDR are plotted relative to normalized enrichment scores (NES). Circle size denotes the number of genes in each category and circle color indicates FDR q values (E-F) Heat map of normalized DESeq2 counts (E) and mRNA levels by qPCR (F) for disallowed genes in islets isolated from 12w old mice p<0.05 (G-H) Heat map of normalized DESeq2 counts (G) and mRNA levels by qPCR (H) for pancreatic progenitor genes in islets isolated from 12w old mice p<0.05. n=6-8 for qPCR analyses. Unpaired Student’s t-test unless stated otherwise. Data are means ± SEM, *p<0.05, ***p < 0.001.

### Changes in chromatin accessibility at β-cell-specific genomic enhancers in dedifferentiated β-cells

ATAC-seq in islet isolated from Tg7 mice revealed that a similar number of accessible regions were detected in 2w (n=63848) and 12w islets (n=65111). A total of 539 and 9840 differential accessible regions (DARs) were detected in 2w and 12w Tg7 mice, respectively (Figure 4a-b, Supplementary Table 2). Intersection of both ATAC-seq datasets revealed that 82.3% (444 of 539) of DARs in non-diabetic 2w mice still remained accessible in hyperglycemic 12w mice, indicating stable remodeling of chromatin landscape following β-cell dedifferentiation. We then assigned regions into one of three categories: opened, closed and unchanged. We observed that the vast majority of accessible chromatin regions in Wt islets of both 2w and 12w old mice failed to open in Tg7 islets (450/539 in 2w mice and 6432/9840 in 12w old mice)(Figure 4c). These DARs were located preferentially in intergenic regions and associated to genes known to regulate the differentiation state of β-cells (*INS1*, *NEUROD1*, *RFX6*, *SYT4*, *SLC2A2*) (Figure 4d-f and Supplementary Figure 3a-c) or located in the vicinity of diabetes susceptibility genes identified in GWAS (*MTNR1B*, and *G6PC2*)(Supplementary Figure 3d-f). Furthermore, chromatin regions failing to open in 2w Tg7 mice were enriched for the presence of DNA motifs for β-TFs, such as Nkx6-1, NeuroD1, MafA/B and Foxa2 (Figure 4g), suggesting that remodeling of chromatin landscapes impairs β-TF recruitment and transactivation of β-cell-specific genes prior to the onset of hyperglycemia. Interestingly, we found that β-cell dedifferentiation predominantly affected chromatin accessibility at enhancer regions decorated by activating H3K4me1^+^/H3K27Ac^+^ known to control genes maintaining β-cell identity and identified in GWAS studies (Figure 4g and Supplementary Figure 3)(40). Concordantly, the expression of genes associated with these differentially closed regions was significantly lower in mutant islets than all other expressed genes (Figure 4i-j). Together, these results indicate that the miR-7-mediated dedifferentiation decreases chromatin accessibility at enhancers of β-cell regulatory genes.

**Figure 4.**
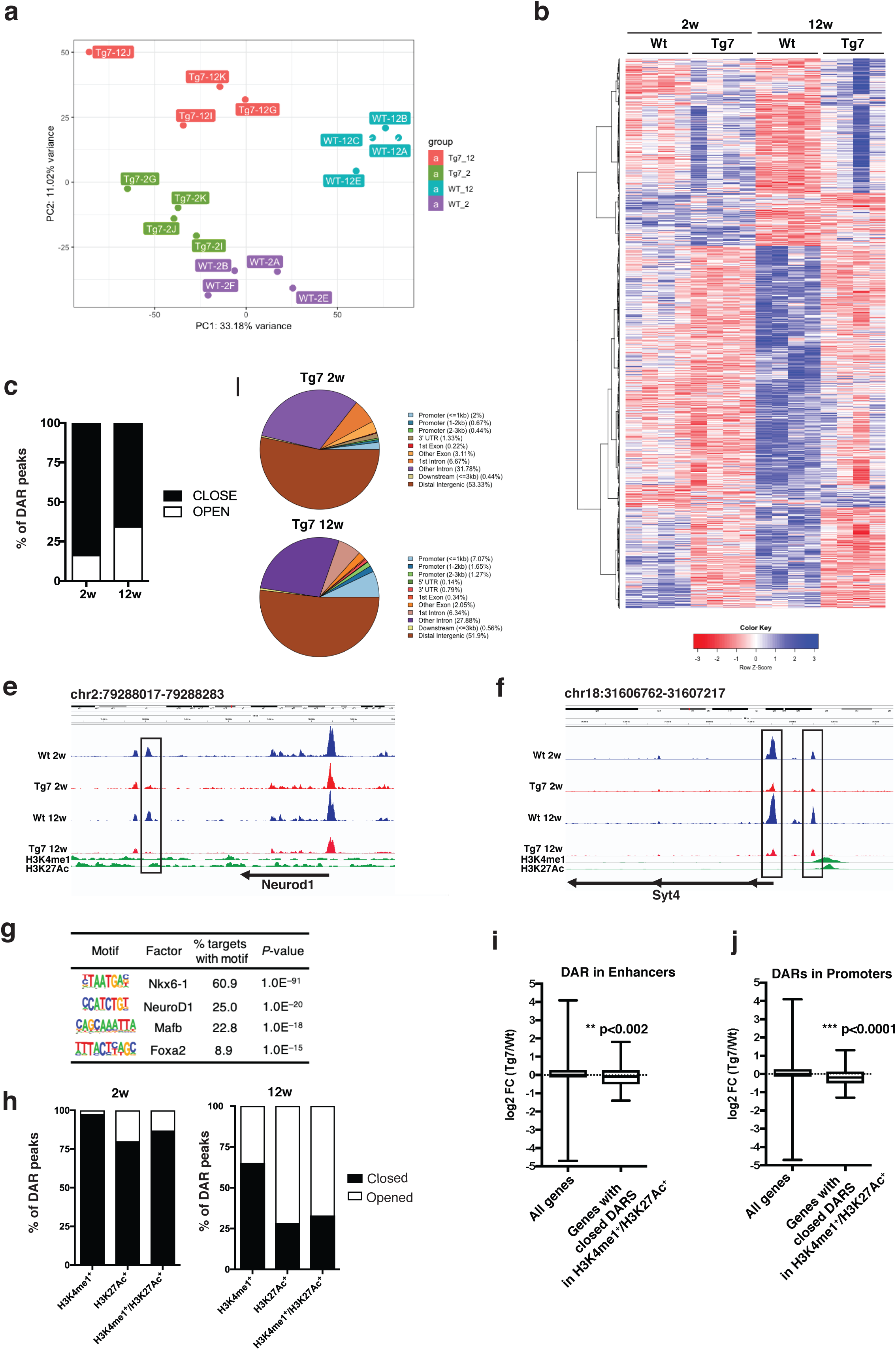
Alteration of chromatin landscapes in dedifferentiated β-cells. (A) Principal component plot (PCA) plot of the ATAC-seq data obtained in islets isolated from 2 and 12-week-old Wt (Wt-2w and Wt-12w) and Tg7 (Tg7-2w and Tg7-12w) mice. Each dot represents a sample (B) Heat map of differentially differential accessible regions (DARs) detected in the ATAC-seq from islets isolated from Wt and Tg mice. padj <0.05 (C) Percentage of islet DARs opening or closing in Tg7 mice padj <0.05 (D) Genomic annotation of DARs in 2w and 12w Tg7 mice (padj <0.05) (E-F) Representative screen shots depicting closed DARs associated to β-cell identity genes (Neurod1 and Syt4) in Tg7 mice. Data processed in IGV 2.4.11 (G) Motif enrichment in closed DAR of Tg7 mice ordered by p-value. (H) Percentage of islet DARs regions decorated with indicated histone 3 modification in Tg7 mice padj <0.05 (I-J) Box plot depicting the expression of genes with closed DARs in H3K4me1^+^/H3K27Ac^+^ enhancer (I) or promoter regions in 12w old Tg7 mice. Unpaired Student’s t-test used unless stated otherwise.

### β-cell dedifferentiation is associated with loss of epithelial identity

Genomic Regions Enrichment of Annotations Tool (GREAT)(41) analyses revealed that ATAC-seq DARs detected in Tg7 mice were enriched in genes involved in epithelial cell functions such as cell adhesion, integrin signaling, cytoskeletal rearrangement and migration (Supplementary Table 3, Supplementary Figure 3g-h), suggesting that β-cells undergoing dedifferentiation concomitantly lose their epithelial identity. In fact, a validation phase on independent islets samples revealed that the expression of a number of epithelial and polarity genes, including *E-cadherin*/*CDH1*, *CLDN4*, *CELSR1*, *PARD6G*, *ACTN4*, *ROBO1* and *TJP2,* was decreased in Tg7 islets (Figure 5a). Further analysis of our RNA-seq data revealed that the top categories of upregulated genes in both 2w and 12w Tg7 islets were also associated with *Epithelial to mesenchymal transition* (EMT) as well as *Inflammation*, *K-Ras activation*, *Myofibroblast activation*, *TGFβ signaling* and *Cell Polarity* (Figure 5b-c). This strongly suggests that β-cell dedifferentiation is associated with loss of the epithelial identity leading to the induction of an EMT process. We confirmed increased expression of EMT markers N-Cadherin (Cdh2), Fn1 and Vim in Tg7 islets as well as Snail1 and Zeb2, two transcription factors mediating EMT signaling (Figure 5d-f and Supplementary Figure 4a)(42). Consistent with TGFβ playing a crucial role in driving EMT, we measured increased levels of p-Smad3 (Figure 5g) as well as induction of several TGFβ responsive genes in Tg7 islets (Figure 5h-i). Interestingly, β-cells from Tg7 islets showed loosened cell:cell contacts and changed their shape from cuboidal to flattened and elongated (Figure 2c and Figure 5j-k). Furthermore, we observed induction of *bona fide* mesenchymal markers (*THY1*, *VCAM1* and *PDGFRβ)* and stemness genes (*N-MYC*, *c-KIT*, *LIF*, *ALDHA2* and *CD9*) in islets of mutant mice (Supplementary Figure 4b-d). Expression of Mef2c, *α*-Sma and Des was increased in Tg7 islets as well as that of collagen genes and ECM remodelers (*MMP2/14* and *SERPINE2*), consistent with the activation of pathways promoting islet fibrosis (Supplementary Figure 4e-i). Given that activated stellate cells represent a major source of TGFβ ligands and proteins regulating matrix secretion and turnover (e.g. Mmps, Timps, Collagens) (43, 44), our data suggests that these myofibroblasts may play an important role in driving this EMT process.

**Figure 5.**
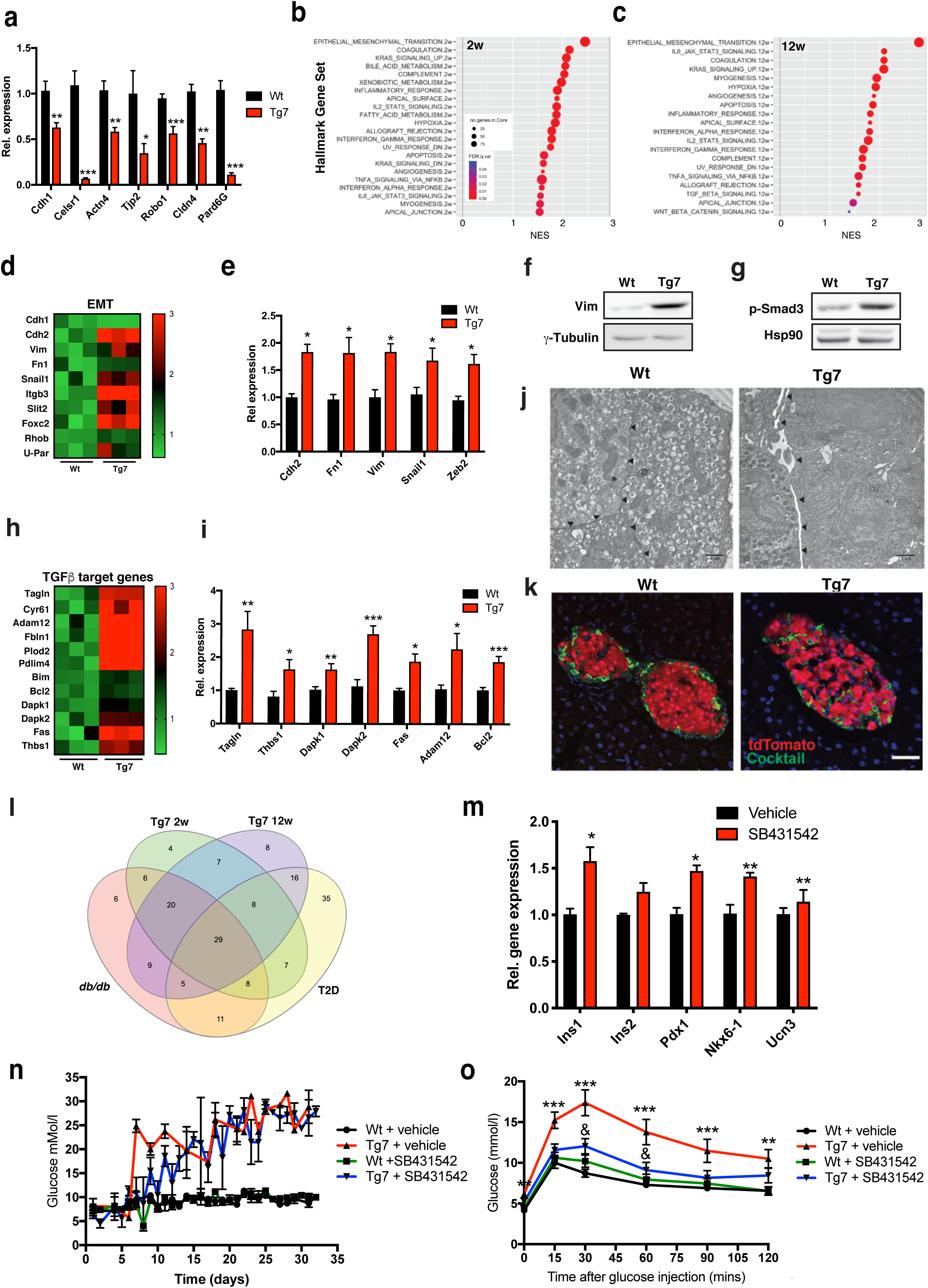
Dedifferentiation of β-cells is associated with loss of epithelial identity and induction of EMT in mouse and humans. (A) Expression of epithelial-specific genes in islets from 12-week-old (12w) Tg7 mice. (B-C) Gene set enrichment analysis (GSEA) of MSigDB Hallmark genes in upregulated pre-ranked gene ratios (Tg:Wt) from 2w (B) and 12w (C) islets. Gene sets with indicated FDR are plotted relative to normalized enrichment scores (NES). Circle size denotes the number of genes in each category and circle color indicates FDR q values (D) Heat map of normalized DESeq2 counts for indicated EMT genes in Tg7 islets versus Wt controls, p<0.05 (E) Expression of EMT genes in islets from 12w Tg7 mice (n=6) (F-G) Western blotting for Vimentin (Vim)(F) and p-SMAD3 (G) in 12w Tg7 islets normalized for indicated loading controls. (H) Heat map of normalized DESeq2 counts for indicated TGFβ target genes in Tg7 islets versus Wt controls, padj<0.05 (I) Expression of TGFβ target genes in islets from 12w Tg7 mice (n=6) (J) Electron microscopy of 12w Wt and Tg7 islet preparations. Arrowheads indicate the intercellular space between β-cells in Tg7 compared to controls islets. Scale bar: 1μm (K) Immunofluorescence with a cocktail of non-β hormone (glucagon, somatostatin, ppy - green) on pancreatic sections from Wt^tdTomato^ and Tg7^tdTomato^ mice. Red: tdTomato autofluorescence. Scale bar: 40μm. (L) Venn diagram of differentially expressed genes from the MSigDB Hallmark *Epithelial Mesenchymal Transition* gene signature in islets from 2w and 12 w Tg7 versus 14-week old *db/db* mice and hT2D datasets (47); padj <0.05 (M) Adult mouse islets were treated with SB431542 for 72h and expression of indicated genes measured by qPCR (n=4) (N) *Ad-libitum*-fed blood glucose levels in Wt and Tg7 mice injected with SB431542 (5mg/kg) three time per week from 2 week-of-age (n=4-17/genotype) (O) IPGTT (0.25g/kg) in overnight fasted Tg7 and control mice at 7 weeks of age (34 days of treatment);(n=6-16/genotype); two-way ANOVA analysis. *p<0.05, **p<0.01 and ***p<0.001, In M, ** (p<0.01), *** (p<0.001) relates to Wt vs Tg7, whereas & (p<0.05) relates to Tg7 vs Tg7+SB431542

To explore the significance of this EMT signature in diabetes, we analyzed the transcriptome of *db/db* islets, a diabetic mouse model displaying increased miR-7 activity and β-cell dedifferentiation (32, 45, 46). Downregulated genes in *db/db* islets were associated with the gene set *Pancreatic β-Cells*, whereas genes upregulated were predominantly associated with an *EMT* signature (Supplementary Figure 5a-b), which correlated with increased number of Ins^+^Vim^+^ β-cells Supplementary Figure 5c). More remarkably, we found that individuals with T2D also displayed enrichment for genes associated with EMT (Supplementary Figure 5d-e)(47), revealing the significance of EMT in humans. Interestingly, hyperglycemia did not appear to contribute significantly to the induction of EMT genes as revealed by the similar overlap between the EMT gene cluster in normoglycemic Tg7 2w (n=91) versus diabetic Tg7 12w (n=86) and *db/db* islets (Figure 5l), supporting the view that the EMT process is an intrinsic β-cell autonomous process. Importantly, we identified 29 EMT-associated genes enriched in Tg7 and *db/db* mice and islets from T2D patients (Figure 5l and Supplementary Table 4). These results indicate that an EMT is a key process that accompanies the dedifferentiation of β-cells in T2D.

To test how EMT affects β-cell identity and function, we first treated islets with SB431542, an anti-EMT drug and inhibitor of TGFβR-I (48). QPCR revealed that SB431542 increased mRNA levels of Ins1, Pdx1, Nkx6-1 and Ucn3 (Figure 5m), indicating that TGFβ signaling is constitutively active in healthy β-cells and limits the expression of key β-cell-specific genes. To test the pathological significance of this EMT process in diabetes, SB431542 was injected in normoglycemic Tg7 and control mice from 2-week of age and glycemia was measured over time. We observed that Tg7 mice injected with SB431542 tend to show a delay in the onset of hyperglycemia (Figure 5n). More interestingly, treatment of mice with SB431542 completely normalized the glucose intolerance of fasted Tg7 mice (Figure 5o), thus revealing that EMT promotes β-cell dysfunction in diabetes.

### β-TFs regulate an epithelial and mesenchymal gene program

Since we identified a strict correlation between β-cell dedifferentiation and EMT, we hypothesized that loss of β-TFs expression my underly loss of epithelial identity and mesenchymal reprogramming of β-cells through EMT. Actually, mining of publicly available mouse islet ChIP-seq datasets for the β-TFs Pdx1, Neurod1 and Nkx6-1 (40, 49) revealed that the genomic loci of several epithelial and mesenchymal markers are bound by β-TFs and appear to be differentially regulated in islets from 2w and 12w Tg7 as well as *db/db* mice (Figure 6a-b). To determine whether *β−*TFs modulate the expression of these genes, we depleted the expression of Pdx1, Nkx6-1 and Neurod1 in MIN6 cells using RNA interference (RNAi) and quantified the expression of epithelial and mesenchymal genes. Inactivation of β-TFs resulted in a marked decreased in epithelial gene expression and concomitantly increased the expression of genes previously associated with mesenchymal reprograming (Figure 6c-h). Knockdown of any of the three β-TFs resulted in decreased expression of Ehf, an epithelial specific transcriptional regulator whereas inactivation of Pdx1 and Nkx6-1 impaired the expression of tight junction protein Tjp2 as well as Celsr1, a planar cell polarity gene driving β-cell differentiation during development (50)(Figure 6c-e). On the other hand, β-TF inactivation correlated with increased expression of known mesenchymal markers such as Snail2, Serpine2, Rgs4, CD44, Pdlim4 and Cyr61 (Figure 6f-h). Amongst these genes, we could validate that Pdx1 is directly recruited to CD44 loci a cell-surface adhesion glycoprotein involved in the progression of EMT (Figure 6i), providing further evidence for direct regulation of mesenchymal genes by β-TFs. Together, these results indicate that activation of epithelial genes and continual repression of mesenchymal genes by β-TFs is required to maintain the functional and epithelial identity of β-cells.

**Figure 6.**
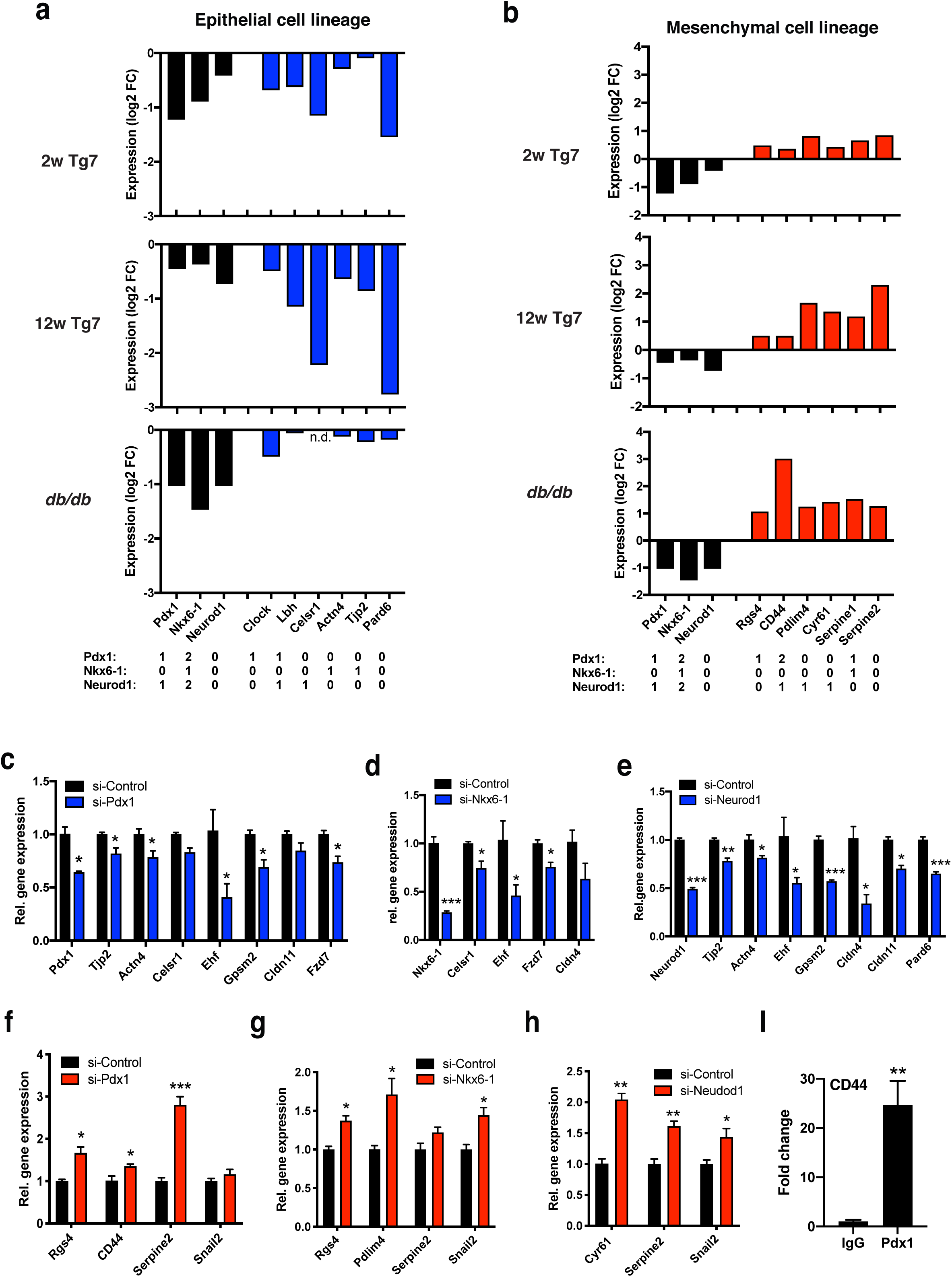
Regulation of epithelial and mesenchymal lineage genes by β-TF in β-cells. (A-B) Log2 fold change expression of β-TFs as well as epithelial and mesenchymal cell lineage genes obtained from RNA-seq of islets isolated from Tg7 (2 and 12 week of age) and *db/db* mice (45); padj <0.05 (C-E) Expression of epithelial-specific genes targeted by β-TFs in MIN6 cells depleted of Pdx1 (C), Nkx6-1 (D) or Neurod1 (E) by RNAi (n=3) (F-H) Expression of mesenchymal-specific genes targeted by β-TFs in MIN6 cells depleted of Pdx1 (F), Nkx6-1 (G) or Neurod1 (H) by RNAi (n=3). Serpine2 and Snail2 do not have β-TF binding sites and are indicative of mesenchymal reprogramming. (I) ChIP for Pdx1 in MIN6 cells demonstrating direct recruitment of the transcription factor to the mesenchymal CD44 gene (n=30). Unpaired Student’s t-test used unless stated otherwise. *p<0.05, **p<0.01 and ***p<0.001

### Smarca4/Brg1 is a target of miR-7 in β-cells

To identify direct miR-7 targets that impair the functional and epithelial identity of β-cells, we probed our RNA-seq data for predicted miR-7 targets genes. This revealed that that islets from 2w and 12w Tg7 mice were enriched for downregulated genes that possess evolutionary conserved miR-7 binding sites in their 3-UTR (padj. 2.13e10^-7^ at 2w and 2.33e10^-9^ at 12w)(Supplementary Figure 6a-b and Supplementary Table 5), with >77% of predicted target genes being downregulated in islets of both 2w and 12w Tg7 mice (Figure 7a). Amongst these genes was *SMARCA4 (BRG1*), which encodes a catalytic subunit of the mammalian Switch/Sucrose Non-Fermentable (mSwi/Snf) nucleosome remodelling complexes (Figure 7b and Supplementary Table 5). Interestingly, mSwi/Snf complexes are key regulators interacting with lineage defining transcription factors controlling both cellular identity (51–54) and inactivating mutations in Brg1 promotes epithelial cell reprogramming and EMT (55, 56). Yet, very little is known on the biological function of Brg1 in pancreatic β-cells and diabetes. Reporter assays performed in cells transfected with a reporter bearing the Brg1 3-UTR cloned downstream of a luciferase gene revealed that miR-7 mimics significantly reduced luciferase activity compared to control transfected cells (Figure 7c). Concordant with Brg1 mRNA being repressed by miR-7, we measured decreased Brg1 mRNA and protein levels in β-cells of Tg7 mice (Figure 7d-e). Similarly, infection of dissociated mouse and human islets with Ad-miR-7 also decreased Brg1 protein levels (Figure 7f-g). Together, these results indicated that Brg1 is a direct target of miR-7 in β-cells.

**Figure 7.**
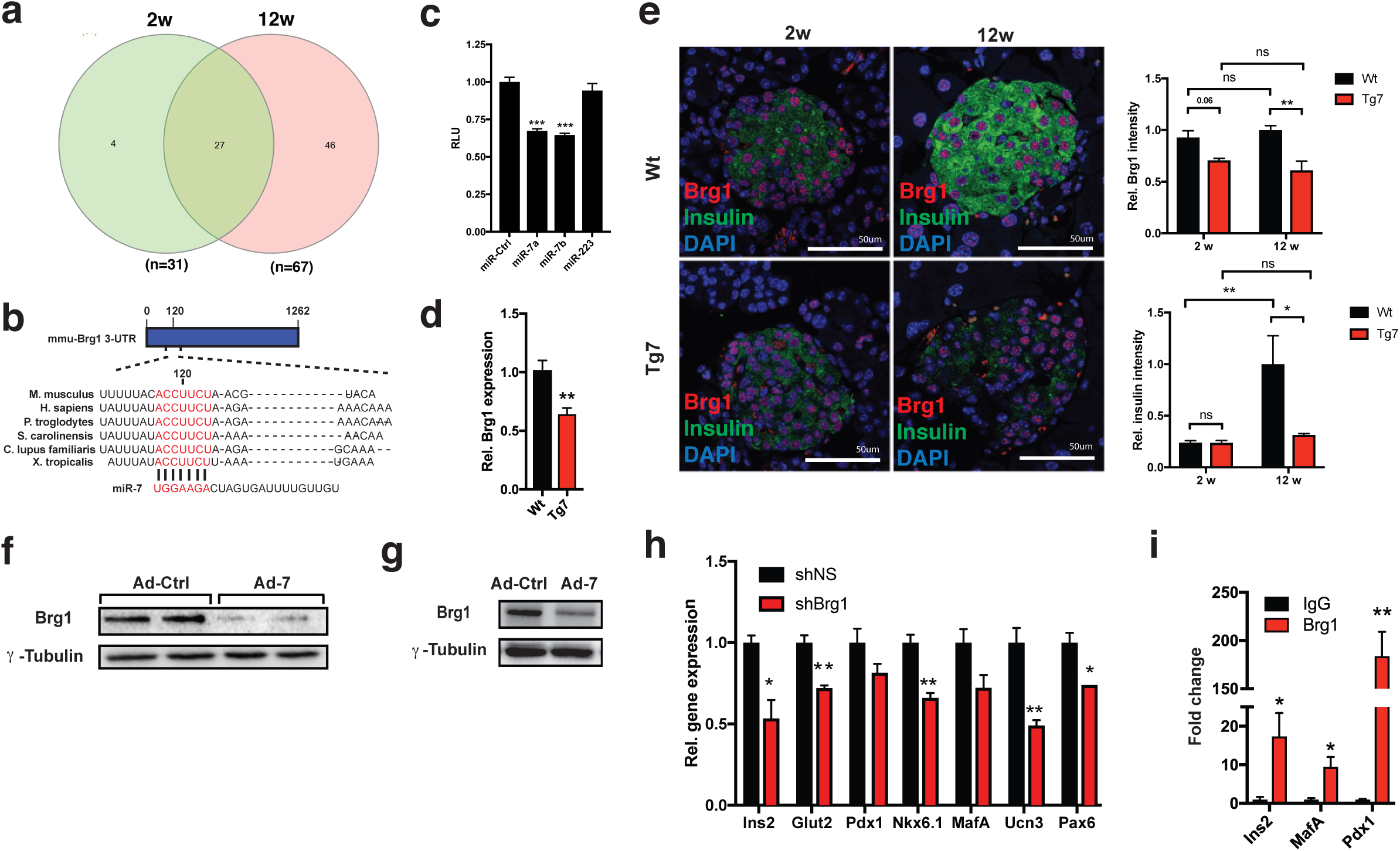
miR-7 represses Brg1 expression in pancreatic β-cell. (A) Venn diagram of predicted miR-7 target genes (Target scan) found to be downregulated and possessing a miR-7 binding site in their 3-UTR in islets isolated from both 2-(2w) and 12-week-old (12w) Tg7 versus Wt mice, padj<0.05 (B) Evolutionary conserved miR-7 binding-site in the 3-UTR of Brg1 mRNA. Nucleotide numbering based on the sequence found in mouse (*Mus musculus*) (C) Relative luciferase levels of a construct harbouring the 3’UTR of Brg1 in HEK293 cells cotransfected with miR-7a, −7b or non-targeting controls (miR-Ctrl and miR-223); n=3, Two-way ANOVA (D) Brg1 mRNA expression in islets isolated from 12w Tg7 mice (n=7) (E) Insulin and Brg1 co-immunofluorescence of pancreatic sections from Wt and Tg7 mice at 2w and of age. Right: corresponding quantification; n=3/genotype. Two-way ANOVA (D-E). (F-G) Western blotting analysis of Brg1 in dissociated mouse (F) and human (G) islets infected with Ad-Ctrl or Ad-miR-7 adenovirus (H) mRNA levels of indicated β-cell genes in INS1E cells stably expressing Brg1 shRNA (shBRg1) or non-silencing shRNAs (shNS), n=3. (I) MIN6 ChIP-qPCR analysis for Brg1 recruitment at indicated loci. n=3 per condition. Unpaired Student’s t-test used unless stated otherwise. Data are means ± SEM, *p < 0.05, **p < 0.01, ***p < 0.001

### Genetic inactivation of Brg1 causes β-cell dedifferentiation, loss of epithelial identity and mesenchymal reprogramming

We then investigated the role of Brg1 in maintaining the identity of mature β-cells. Loss-of-function experiments in MIN6 cells revealed that downregulation of Brg1 by RNA interference (RNAi) or CRISPR/Cas9 gene editing resulted in decreased expression of a number of genes involved in preserving β-cell identity (Figure 7h, Supplementary Figure 6b-c). Chromatin immunoprecipitation qPCR (ChIP-qPCR) analysis demonstrated that Brg1 is recruited at genomic loci encoding β-cell specific genes including *INS2*, *MAFA* and *PDX1* (Figure 6I). Not only do these results demonstrate that Brg1-containing complexes are required to maintain the expression of β-cell identity genes, but they also provide a molecular mechanism for the overall compaction of chromatin landscapes in dedifferentiated β-cells (Figure 4).

To reveal the importance of Brg1 in preserving the functional and epithelial identity of β-cells *in vivo*, we conditionally inactivated Brg1 in β-cells using mice with a floxed Smarca4/Brg1 gene and bearing a Ins1-Cre transgene (Brg1*^f/f;Ins1Cre^*)(57). Quantitative analysis revealed a 70% downregulation of Brg1 expression in islets from mutant mice (Figure 8a). Examination of controls and mutant pancreata did not reveal any changes in islet morphology (Figure 8b). However, we observed an overall decrease in insulin protein expression in β-cells of Brg1*^f/f;Ins1Cre^* mice compared to controls (Figure 8b). This also correlated with a significant reduction of several β-cell differentiation markers including *INS1/2*, *MAFA* and *SYT4* (Figure 8c)(58). Moreover, Brg1 expression is required to maintain the epithelial identity of β-cells as the expression of several epithelial markers was markedly decreased in mutant islets (Figure 8d). Furthermore, we could detect increased expression of a number of EMT-related genes including Serpine2, Rgs4, CD44 in islets from Brg1*^f/f;Ins1Cre^* mice (Figure 8e). These results support that miR-7 impairs the functional and epithelial identity of β-cells via repression of Brg1 expression.

**Figure 8.**
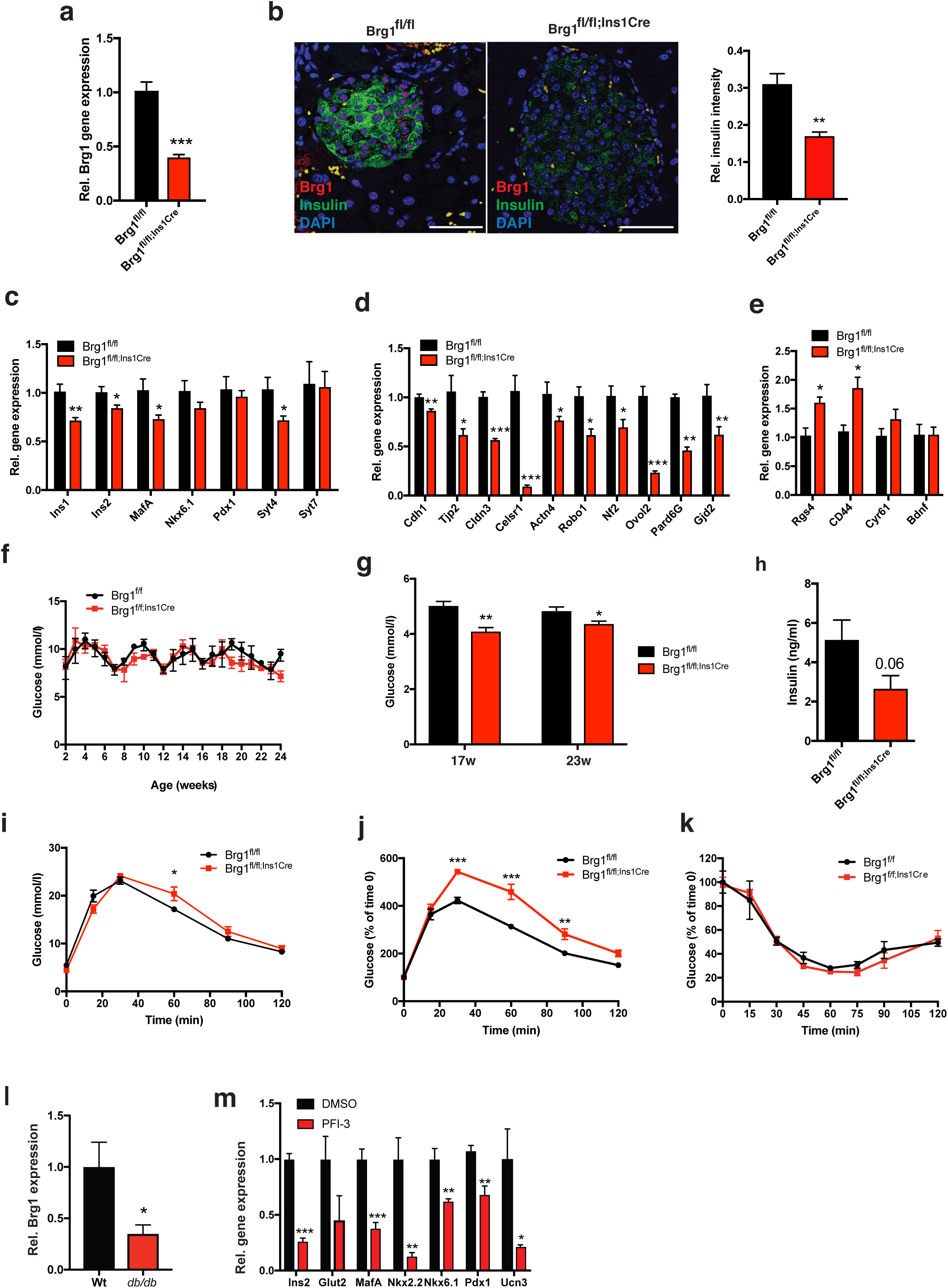
Impaired β-cell and epithelial identity in conditional Brg1 knockout mice. (A) Brg1 expression in islets isolated from 24-week-old Brg1^fl/fl;Ins1Cre^ and control (Brg1^fl/fl^) mice (B) Insulin and Brg1 co-immunofluorescence on pancreatic sections of Brg1^fl/fl;Ins1Cre^ and control mice. Right: Quantification of insulin immunostaining (n=3-4/group) Scale bar: 50 μm (C-E) mRNA levels of indicated β-cell-specific genes (C), epithelial (D) and mesenchymal (E) markers in islets from Brg1^fl/fl;Ins1Cre^ and controls (n=6/genotype) (F) *Ad-libitum*-fed blood glucose levels in Brg1^fl/fl;Ins1Cre^ and control mice (n=8-10/genotype) (G) Glycemia of overnight fasted Brg1 ^fl/fl;Ins1Cre^ mutant and control mice of indicated age (n=7 /group) (H) Circulating insulin levels in 15-week-old Brg1^fl/fl;Ins1-Cre^ and control mice (K) mRNA levels of Brg1 (n=7/group) (I) IPGTT (2g/kg) in overnight fasted Brg1^fl/fl;Ins1Cre^ and control mice at 16 weeks of age; (n=8-10/genotype); two-way ANOVA analysis (J) Data from panel I normalized glycemia before glucose injection. two-way ANOVA analysis (K) IPITT (0.75U/kg) in 5-hour fasted Brg1^fl/fl;Ins1Cre^ and control mice at 17 weeks of age; (n=8-10/genotype); Two-way ANOVA (L) Expression of Brg1 in islets from 14-week-old *db/db* mice and controls (n=5). (M) Expression indicated β-cell specific genes in human islets treated with the Brg1 inhibitor PFI-3 (n=4). Unpaired Student’s t-test used unless stated otherwise. Data are means ± SEM, *p < 0.05, **p < 0.01, ***p < 0.001

Metabolic characterization of Brg1*^f/f;Ins1Cre^* mice revealed no significant change in random-fed glycemia (Figure 7F). Overnight fasting resulted in decreased circulating glucose levels (Figure 8g), suggesting impairment of counter-regulatory mechanisms of glycemic control from islet cells. Supporting loss of β-cell identity, we found lower circulating insulin levels in mutant versus Wt (Figure 8h). When challenged with an intra-peritoneal glucose tolerance test (IPGTT), we observed a mild glucose intolerance in Brg1*^f/f;Ins1Cre^* mice versus controls (Figure 8i), which may be attributed to decreased fasting glycemia in Brg1*^f/f;Ins1Cre^* mice before injection of glucose. However, when the data is normalized for the glycaemia of each animal at t=0, Brg1*^f/f;Ins1Cre^* displayed marked glucose intolerance, while presenting no change in insulin sensitivity (Figure 8j-k), thus supporting a β-cell defect in Brg1 animals. Importantly, we found that Brg1 mRNA levels are decreased by over 50% in *db/db* islets, supporting its role in maintaining β-cell identity (Figure 8l). Finally, to provide evidence for a role of Brg1 in maintaining the identity of humans β-cells, we pharmacologically inhibited Brg1 in human islets using PFI-3, a selective bromodomain inhibitor with high affinity for Brg1 (59). Treatment with PFI-3 resulted in downregulation of several β-cell identity genes (*INS*, *MAFA*, *NKX6.1*, *PDX1* and *UCN3)* compared to controls (Figure 8m). Together, our results indicate that miR-7-mediated regulation of Brg1 represents a core component of an evolutionarily conserved circuit stabilizing the functional and epithelial identity of pancreatic β-cells.

## DISCUSSION

Pancreatic β-cells originate from epithelial precursor cells and their polarity is defined by surface microdomains composed of distinct cell adhesion molecules, cell:cell contacts (tight, gap and adherens junctions) and signaling molecules that are required for β-cell function (60–64). Compromised epithelial cell identity results in the induction of an EMT process and underlies the development of many diseases (65–67). Here, we uncover that β-cell dedifferentiation is associated with an EMT process reminiscent of a response to tissue injury which is mediated by miR-7-mediated repression of Brg1, a catalytic subunit of the Swi/Snf chromatin remodeling complexes. We found that Brg1 is required to preserve chromatin accessibility at H3K4me1^+^/H3K27Ac^+^ decorated lineage defining enhancers controlling the expression of β-cell specific and epithelial genes in mature β-cells (Figure 2-5)(68, 69). Failure to maintain this epigenetic regulatory program in diabetes leads to decreased expression of β-cell specific genes and markers of epithelial cells as well as induction of genes involved in mesenchymal reprogramming. Indeed, we identify 29 EMT-associated diabetes therapeutic target genes, most of which have not been previously reported to be associated with β-cell dysfunction in both mouse and humans. More remarkably, pharmacological inhibition of this EMT process improves pancreatic β-cell function of diabetic mice, further highlighting the pathological relevance of EMT in T2D. Despite loosened β-cell:β-cell contacts prompted by β-cell dedifferentiation and EMT, we did not observe pancreatic tumor nor dissemination of Vim^+^ dedifferentiated β-cells in any diabetic mouse model (data not shown), indicating an EMT process leading to fibrosis as opposed to migratory phenotype associated with metastasis.

In search for the molecular mechanism underlying activation of EMT in dedifferentiated β-cells, we discovered that β-TFs are fundamental guardians of β-cell identity through their ability to activate and repress genes defining epithelial and mesenchymal cell fate, respectively (Figure 6). Our work expands the repertoire of non-β-cell genes continually repressed by β-TFs by unexpectedly revealing that, in addition to repress hormone genes such as somatostatin or glucagon (12, 14, 15, 17, 22), β-TFs inhibit mesenchymal lineage-defining genes. In light of the interaction of the mSwi/Snf complex with Pdx1 and Nkx6-1, (70–72), we propose that increased miR-7 levels in T2D islets downregulates Brg1 expression and triggers β-cell dedifferentiation by affecting, on one hand, the activation of genes encoding β-TFs and regulators involved in glucose sensing, membrane excitability, metabolism, but on the other hand, epithelial genes involved in regulation of cell:cell contact, communication and polarity. As a direct consequence of lowered concentration of β-TFs, de-repression of a subset of mesenchymal genes is elicited and an EMT process is induced. As we are witnessing a growing body of evidence indicating induction of the EMT marker Vim in β-cells of diabetic subjects (73–76), our data provide mechanistic insights into this EMT process in T2D.

Previous studies have shown that cellular dedifferentiation represents a key step in the regeneration of damaged or stressed cells in several organs (77) and is associated with extensive alteration of the tissue environment leading tin some case to fibrosis. For example, an EMT-like process is activated following peripheral nerve injury and underlies cellular reprogramming of Schwann cells which drive nerve regeneration (78). However, genetic lineage studies in the pancreas indicate that reprogramming of β-cell identity in response to metabolic stress appears to underlie a pathological, maladaptive mechanism in diabetes rather than a regeneration process leading to recovery of the β-cell mass. However, in extreme diabetes mouse models with β-cell depletion, regeneration of insulin-producing cells has been shown to be controlled by TGFβ signaling (79), whereas acute β-cell depletion by streptozotocin triggers the repopulation of β-cell mass from a rare population of Vim^+^MafB^+^ cells (80). Given that EMT is a multi-stage process (81, 82), one possibility is that the EMT process we report here represents a snap shot of an intermediary stage of a long-term regenerative process aimed at restoring β-cell mass.

In summary, our study unveils a novel post-transcriptional mechanism regulated by miR-7 in β-cells. In addition to the well documented role of Brg1 in cancer (83), we reveal that Brg1 is at the center a crucial regulatory mechanism maintaining pancreatic β-cell fate and, in turn, have important implications for mSwi/Snf complexes in driving EMT in metabolic disease. As we are witnessing the first Phase II clinical programs testing the efficacy of antagomirs blocking the function of a specific miRNA in different types of disorders (84, 85), we believe that in addition to anti-EMT drugs, miR-7 inhibitors may represent innovative ways to foster the functional and epithelial identity of β-cells or prevents their mesenchymal reprogramming in T2D.

## METHODS

### Animal Housing conditions

All animals were kept and bred in a pathogen-free animal facility according to the Home Office regulations as defined by the Animal (Scientific Procedures) Act, 1986. Mice were housed in Allentown XJ individually ventilated cages (IVCs) on a 12-hour light/dark cycle with constant environmental conditions (temperature: 21°C ± 2°C, humidity: 55% ± 10%) and had free access to standard rodent chow. All procedures were performed in accordance with UK Home Office regulations under HO Project Licence number 70/8967 (M. Latreille).

### Animals

Tg7 mice have been previously described (32). Floxed Brg1 mice (kindly provided by Dr Pierre Chambon via M. Goetz, Helmholtz Institute, Munich) were crossed with Ins1-Cre mice (57) to generate the β-cell specific Brg1*^f/f;Ins1-Cre^* line. Genetic lineage tracing was performed following crossing of Tg7 mice with rat Ins2 (RIP)-Cre mice (86) carrying a Rosa26-floxed-Stop-tdTomato transgene. Glucose and insulin tolerance tests were performed in fasted mice (16h hr for IPGTT and 5h for IPITT). D-glucose was injected intraperitoneally at 2g/kg or 0.25g/kg body weight as indicated. Insulin was injected intraperitoneally at 0.75 U/kg body weight. Blood glucose levels were measured by tail venesection at the indicated times. SB421542 (Sigma) was dissolved in 50% DMSO and injected intraperitoneally three time per week at 5mg/kg. Male mice (C57BL/6 background) were used in all experiments

### Histology and immunofluorescence

At least three sections ∼200μm apart from three animals of each genotype were used for every analysis. Sections were stained with hematoxylin and eosin as previously described (Fischer et al., 2014). Pancreatic sections were stained using the Picro Sirius Red Stain Kit as described by the manufacturer (Abcam). For immunofluorescence staining, pancreata were dissected, weighted and fixed in 4% PFA at 4°C overnight. Sections were deparaffinized and rinsed in distilled water for 5 min. Antigen retrieval was performed where necessary using a Decloaking Chamber NxGen (BioCare Medical, USA) for 5 min at 110°C for 5 min in Tris-HCl pH 10.0 or sodium citrate pH 6.0. Sections were permeabilized for 30 min in a permeabilization buffer (0.1% Triton/PBS) and blocked for 1 hour at room temperature in blocking buffer (1% bovine serum albumin/5% serum/PBS). Cryosections were directly permeabilized for 15 min in permeabilization buffer and blocked for 1 hour at room temperature in a blocking buffer. Primary antibodies were diluted in blocking buffer and incubate on sections overnight at 4°C. Slides were washed three times for 15 min in permeabilization buffer. Sections were then incubated with a fluorochrome-conjugated secondary antibody diluted in blocking buffer for 1 hour at RT in the dark. Slides were rinsed and mounted on glass slides with VectaShield. Dissociated islets were fixed in 4% PFA/PBS for 15min at room temperature, followed by 2 washes with 1x PBS. Islet cells were incubated with permeabilization buffer for 15 min at room temperature, followed by three washes with 1x PBS and blocked in blocking buffer for 1 hour at room temperature. Primary and secondary antibodies were added as described above. Images were acquired using Leica SP5II or Olympus IX70 and analyzed using ImageJ software.

### Pancreatic islet studies

Pancreatic islets were isolated following perfusion of the pancreas via the bile duct with 0.2 mg/mL of ice cold liberase (Roche) as described before (32). PFI-3 (Sigma Aldrich) was dissolved in DMSO and added to human islets preparations (Table S6) at 100 nM for 72h at 37°C. For dissociation studies, up to 600 islets were picked into 1.5ml microtube and washed three times with PBS. Islet were then incubated with Accutase at 37°C for 10 min and the reaction was stopped by adding fetal bovine serum (FBS). Islet cells were centrifuged at 1,200 RPM for 5 min and resuspended in RPMI 1640 medium, 11mM glucose supplemented with 10% FBS, 2mM L-glutamine, 100 IU/ml penicillin and 100ug/ml streptomycin and seeded in 24-weell plates on coverslips coated with 804G cell conditioned media (87) at a maximum density of 200,000 cells per well. When embedded, mouse and human islets were fixed for 15 min in 4% PFA. Samples were centrifuged at 3,000 rpm for 3 min and supernatant disposed of. Islets were resuspended in 1ul of 3% UltraPure™ Low Melting Point Agarose (Thermo Fisher) and embedded in OCT compound mounting media (VWR chemicals). 10 uM sections were prepared on a cryostat (Leica). For conventional EM, islets were chemically fixed in 2% paraformaldehyde (EM grade), 2% glutaraldehyde and 3 mM CaCl_2_ in 0.1 M cacodylate buffer for 2 hours at room temperature then left overnight at 4°C in fresh fixative, osmicated, enrobed in agarose plugs, sequentially dehydrated in ethanol and embedded on Epon polymerised overnight at 60°C. Ultrathin 70-nm sections were cut with a diamond knife (DiATOME) in a Leica Ultracut UCT ultramicrotome before examination on a FEI Tecnai G2 Spirit TEM. Images were acquired in a charge-coupled device camera (Eagle) and processed in ImageJ. For calcium imaging, islets were loaded with fluo-8 and imaging were performed essentially as described (34, 88) using a Nipkow spinning disk head. In brief, a solid-state laser (CrystaLaser) controlled by a laser merge module (Spectral Applied Physics) provided wavelengths of 491 nm (rate, 0.5 Hz; exposure time, 600 ms). Emitted light was filtered at 525/50 nm, and images were captured with a 16-bit, 512 × 512 pixel back-illuminated EM-CCD camera (ImageEM 9100-13, Hamamatsu) driven by Volocity^TM^ software (PerkinElmer Life Sciences). For connectivity and correlation analyses, individual β-cell regions of interest (ROIs) were visually identified within each dye-loaded isolated islet that was imaged (typically ∼50 per islet). Mean fluorescence intensity timeplots from each ROI were subjected to correlation analyses for all cell pairs using a custom-made MATLAB script (available upon request). Data were smoothed using a retrospective averaging method (previous 10 values) and all traces were normalised to F0. The Pearson correlation function R between all possible (smoothed) cell pair combinations (excluding the autocorrelation) was assessed and the data were subsequently subjected to a bootstrap resampling to increase the accuracy of the confidence interval of the R statistic, with p<0.001 deemed a statistically significant cell-cell connection. Connectivity data were displayed in two formats. Firstly, the Cartesian co-ordinates of the imaged cells within a given islet were used to construct connectivity line maps. Cell pairs (R>0.25 AND p<0.001 post bootstrap) were connected with a straight line, the colour of which represented the correlation strength (R=0.25-0.5 [green], R=0.5-0.75 [yellow], R=0.75-1.0 [red]). Secondly, Pearson heatmap matrices, indicating individual cell pair connections on each axis (min. = −1; max. = 1) were produced.

### Cell culture, transfection and viral infections

MIN6 cells were cultured in Dulbecco’s Modified Eagle Medium with 4.5 g/L glucose and phenol red (DMEM, Thermo Fisher, UK) supplemented with 15% FBS (Euro-Bio), 1% Penicillin-Streptomycin (Thermo Fisher), 1% Glutamax (Thermo Fisher), 1% sodium pyruvate 100mM (Thermo Fisher) and 0.0005% β-Mercaptoethanol (Sigma Aldrich, UK). INS1E cells were cultured in RPMI, supplemented with 10% FBS, 1% Penicillin-Streptomycin, 1% Glutamax, 1% Pyruvate, 0.0005% β-Mercaptoethanol and 0.2% 1M Hepes pH 7.3 (Thermo Fisher). HEK293T cells were kept in DMEM with 5% FBS, 1% Penicillin-Streptomycin and 1% Glutamax. siRNA (Dharmacon) and were transfected using Lipofectamin 2000 (Thermo Fischer) as recommended by the manufacturer. Cells were harvested between 48 and 72h. HEK293 cells were plated out at a density of 5×10^4^ cells / well in a 24-well plate and transfected with 100 ng of pMirGlo reporter containing mouse Smarca4/Brg1 3-UTR (MmiT075618-MT06 ordered from GeneCopoeia^TM^), 50 nM siRNA mimics for miR-7a, miR-7b or miR-223 as a control. Cells were harvested using the Dual-Glo® Luciferase Assay System (Promega) based on manufacturer indications and analyzed in a FLUOstar Omega filter-based multi-mode microplate reader (BMG Labtech). Adenovirus infections were performed as described previously (32). For lentiviral particle production, HEK293T cells were plated into 6-well plates at 1×10^6^ cells/well and transfected of 1 µg lentiviral vector DNA (pGIPZ encoding shRNAs or plenti-V2-guide RNAs), 100 ng pCMV-VSV-G envelopment plasmid and 900 ng psPAX2 packaging plasmids using Lipofectamine 2000 for 3.5 h at 37°C. Media was then replaced by Min6 media and cells were incubated overnight at 37°C. The next day, media was filtered through 45µm Corning acetate filter and cells were further incubated for 24h in fresh media. 8 µg/mL polybrene (Sigma Aldrich) were added to the filtered media and added on cells. 48h later, puromycin was added at 100 µg/mL and the cells incubated overnight at 37°C. RNA extractions were performed 48h later.

### RNA extraction, quantitative PCR and Western blotting

RNA was extracted using Trizol and treated with DNaseI using RNAse-free DNAse set (Thermo Fisher) based on the manufacturer’s recommendation. Reverse transcription was done using the High-Capacity cDNA Reverse Transcription Kit (Thermo Fisher) and quantitative PCR was performed using KAPA SYBR FAST qPCR Master Mix Kit (Kapa Biosystems) on a LightCycler® 480 apparatus (Roche). Data was normalized over the house keeping gene *RPLP0*. The list of EMT-related genes targeted by β-TFs was determined from the dbEMT gene set (http://dbemt.bioinfo-minzhao.org/). For western blotting, cells lysed in RIPA buffer containing 5% TRIS-HCl pH 8.0, 3% NaCl 5M, 0.4 % EDTA 0.5 M pH 8.0, 10 % NP-40 10%, 10 % Sodium Deoxycholate 10% and proteinase inhibitor in H^2^O (Roche) and samples were treated as previously described (32).

### Statistical Analysis

Data are reported as mean ± standard error of the mean (SEM). Statistical significance was using Student t-test or Anova with Sidak post hoc tests using Prism 7 (GraphPad software) and p<0.05 was considered significant.

## ACKNOWLEDGEMENTS

This work was supported by Medical Research Council Grant MC-A654-5QC20. G.A.R. was supported by a Wellcome Trust Senior Investigator Award (WT098424AIA) and Investigator Award (212625/Z/18/Z), MRC Program grants (MR/R022259/1, MR/J0003042/1, MR/L020149/1) and Experimental Challenge Grant (DIVA, MR/L02036X/1), MRC (MR/N00275X/1), Diabetes UK (BDA/11/0004210, BDA/15/0005275, BDA 16/0005485) and Imperial Confidence in Concept (ICiC) grants, and a Royal Society Wolfson Research Merit Award. Also supported by the European Union’s Innovative Medicines Initiative 2 Joint Undertaking under grant agreement No 115881 (RHAPSODY) to G.A.R. and M.S. This Joint Undertaking receives support from the European Union’s Horizon 2020 research and innovation program and EFPIA. A.T. was supported by Medical Research Council Grant MR/R010676/1. We would like to thank Lorraine Lawrence for histology, Dirk Dormann and Chad Whilding for microscopy and CellProfiler pipeline assistance. We want to acknowledge Ines Cebola for advising on ATAC-seq experiments. We thank members of the Latreille lab, Chris Schiering and Naveenan Navaratnam for insightful discussions and critical reading of the manuscript.

## AUTHOR CONTRIBUTION

T.M., Y.v.O., P.C., and M.L. designed all experiments. T.M., Y.v.O., K. J., P. E., P.C., A.T. and M.L. performed experiments. T.M., Y.v.O., Y. F.W., E. K., K. J., P. E., P.C., W.D., V.S. and M.L. analysed data. M.S. provided mouse lines., P.M., A.M.J.S., G.A.R. coordinated procurement and culture of human islet samples. T.M., Y.v.O, and M.L. prepared the figures. G.A.R wrote a part of the manuscript. M.L. directed the study and wrote the manuscript. All authors read and approved the final version of the manuscript.

## Supplementary information

### SUPPLEMENTARY METHODS

#### Chromatin immunoprecipitation

One 150 mm plate of Min6 cells was washed three times with PBS and incubated with 1% formaldehyde/PBS (Sigma) on a rocking platform for 10 minutes at room temperature. Glycine was added to a final concentration of 0.11 M and incubated on a rocking platform for 5min at room temperature. Cells were washed twice with ice cold 1x PBS on ice, scraped spun down at 2000 g for 2 minutes and aliquoted in 4 tubes containing 5% BSA. Aliquots were washed with 2.5 ml of ice cold 1x PBS with 1x protease inhibitors (Roche). Aliquots were spun down at 2000 g for 2 minutes at room temperature and cell pellets were snap frozen in a dry-ice/ethanol bath and lysed in 140 ul Lysis buffer (2% Triton X-100, 1% SDS, 100 mM NaCl., 10 mM Tris-HCl pH 8.0, 1 mM EDTA with protease inhibitor) and subjected to mechanical disruption using a pestle. The lysed cells were sonicated for 5 minutes (30 sec on / 30 sec off) in a Bioruptor® Pico sonication device (Diagenode). The sonicated samples were spun down at 21000 g for 4 minutes at 4°C and the supernatant was saved in a microtube. 5ul of the sample was used to monitor fragment sizes. For each sonicated sample 110 ul Protein Dyna beads A (Thermo Fisher) was added to the samples. Dyna beads were washed 3x on a magnetic rack with 1ml ice cold Working buffer (1 and 4 parts of Lysis to Dilution buffer (50 mM Hepes (Sigma), 140 mM NaCl, 1mM EDTA, 0.75% Triton X-100, 0.1% Na-deoxycholate (Sigma), 1x PI in H_2_O). Beads were then blocked in 1 ml Preblocking buffer (20 mM glycogen, 0.5% BSA diluted in Working buffer) at 4°C overnight. Beads were washed 3x in 1 ml ice cold Working buffer and resuspended in 100 ul Working buffer. Chromatin was precleared by rotating the sonicated samples with 30ul Dyna beads diluted in Dilution buffer at a 1:4 ratio, in a final volume of 1 ml with Working buffer for 2 hours at 4°C. 50 ul of the supernatant were stored for the Input sample in qPCR. 10 ul of Brg1 or Pdx1 antibody was added to approximately 4 × 10^6^ or 5 ug of antibody to rabbit IgG were added to 30ul of Dyna beads and 1 ml of ice cold Working buffer where 50 ul of 10% BSA and 5 ul Yeast tRNA (Thermo Fisher) were added. Antibody-mixes rotated for 2 hours at 4°C, followed by 3 washes using 1 ml of ice cold Working buffer. Supernatant was disposed and the precleared chromatin was equally divided between Brg1 and IgG antibody-bound beads. Samples were rotated overnight at 4°C. Supernatant were stored as flowthrough for the qPCR analysis. The antibody-coupled beads were pulled down and chromatin fragments washed and rotated in sequence on ice for 5 minutes at 4°C with 1 ml Low salt-(1% Triton X-100, 150 mM NaCl, 20 mM Tris-HCl pH 8.0, 0.1% SDS, 2 mM EDTA), High salt-(500 mM NaCl, 1% Triton X-100, 20mM Tris-HCl pH 8.0, 0.1% SDS, 2 mM EDTA) and LiCl-(0.25 mM LiCl, 1% Sodium deoxycholate, 10 mM Tris-HCl pH 8.0, 1% NP-40, 2 mM EDTA) working buffer, followed by 3 washes with 1 ml TE buffer (10 mM Tris-HCl pH 8.0, 1 mM EDTA). 150 ul of Elution buffer (1% SDS, 0.1 M NaHCO_3_) was added to the beads and samples were put in a thermomixer for 15 minutes at 1200rpm at 65°C. Supernatant were transferred to a fresh microtube, and elution repeated with 150 ul Elution buffer and eluates combined. For DNA isolation 100 ul of the Flow through, the Input, the Fragmentation test aliquot and the eluates diluted to 300ul with TE buffer. 1.5 ul of RNAse A was added to each sample and incubated in a water bath at 65°C for 1h. 4.5 ul of Proteinase K and 12 ul of 5 M NaCl was added and samples incubated in a water bath overnight at 65°C. DNA recovered by adding 60 ul of TE buffer to each samples, before the addition of 450 ul of Phenol:Chloroform. Samples were vortexed and centrifuged for 5 minutes at 21000g. Top phase was recovered and 450 ul of Chloroform were added. Samples were spun down for 5 minutes at 21000g and top phase was kept. Na Acetate (1/10 of the volume) and 2.5 ul of glycogen was added followed by the precipitation in 2.5 volume of ice-cold ethanol. Tubes were inverted and incubated overnight at −20°C. Samples were centrifuged for 20 minutes at 21000 at 4°C and supernatants discarded. Samples were washed with 1 ml of ice cold 70% ethanol, centrifuged and cell pellet was dried and resuspended in a maximum volume of 100ul H_2_O. DNA in H_2_O was incubated in a thermo mixer for 30 minutes at 1000 rpm at 37°C followed by qPCR analysis.

#### RNA-sequencing analysis

RNA was extracted from islets of 2w and 12w Tg7 and littermate Wt controls (n=3, pools from at least 3 animals). RNA-seq libraries were prepared with NEB Ultra II RNA librairy kit (Illumina) from 10 ng total. Sequencing was then proceeded with Hiseq2500 using paired end 100 bp reads at the MRC LMS Genomics facility. Illumina CASAVA 1.8.4 software used for base calling and demultiplexing. Raw RNA-Seq reads were trimmed with trimmomatic to remove adaptors and low-quality reads (v.0.33; Bolger et al., 2014) then aligned against Ensembl Mus musculus genome reference sequence assembly (mm9) and transcript annotations using tophat2 (2.0.11) (1). Gene-based read counts were then obtained using the featureCounts function (2) from Rsubread Bioconductor package (3). Differential expression analysis was performed using DESeq2 Bioconductor package (4). A ranked gene list based on Wald statistics from DESeq2 results and used by GSEA with MSigDB gene sets from ‘H’ collections (5, 6). Stemness gene lists were obtained from the Cancer stem cell database (CSCdb) at http://bioinformatics.ustc.edu.cn/cscdb/. The epithelial gene list was manually generated from literature and web searches.

#### ATAC-sequencing analysis

ATAC-Seq was performed following the instructions as described previously (7). ATAC-seq librairies were prepared using the NEBNext Library kit (Illumina) from Tg7 and Wt control islets (n=4, pools of at least 3 mice) and subjected to sequencing on Hiseq2500 at the MRC LMS Genomics facility. Following demultiplexing step, raw reads were trimmed with Trim Galore! (trim_galore_v0.4.4; https://www.bioinformatics.babraham.ac.uk/projects/trim_galore/). The trimmed reads were then aligned to mm9 using bowtie2 (bowtie2/2.1.0, with –very-sensitive −X 2000) (8). R bioconductor package ATACseqQC (9) was applied for removing duplicated reads and reads mapped to chrMT, then further splitting reads into nucleosome-free (fragment length <=100 bps), mononucleosome, dinucleosome, and trinucleosome bins. We used the nucleosome-free bin for the downstream analysis. Peak calling was proceeded with MACS2 peakcalling function (with -f BAMPE) (10). Peaks overlapped with the blacklist (ENCODE Project Consortium, 2012; https://sites.google.com/site/anshulkundaje/projects/blacklists) were removed and only those appearing in more than any 4 samples were kept. Peak-based read counts were then obtained using the featureCounts function from Rsubread Bioconductor package. Differentially binding analysis was performed with DESeq2 Bioconductor package. Peaks were annotated with ChIPseeker Bioconductor package (11). HOMER (Hypergeometric Optimization of Motif EnRichment) *findMotifsGenome.pl* (with -size 200 -mask) was applied for finding enriched motifs. Histones peaks were from obtained from the following studies:

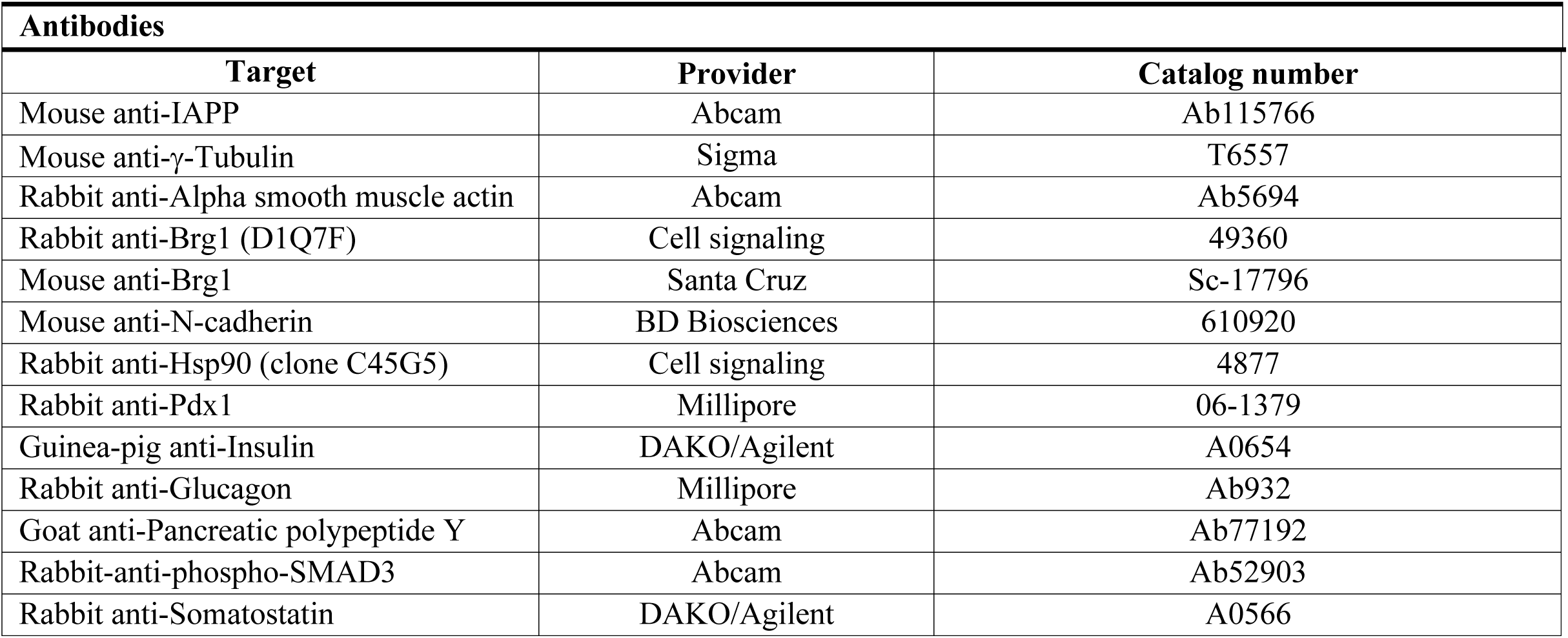
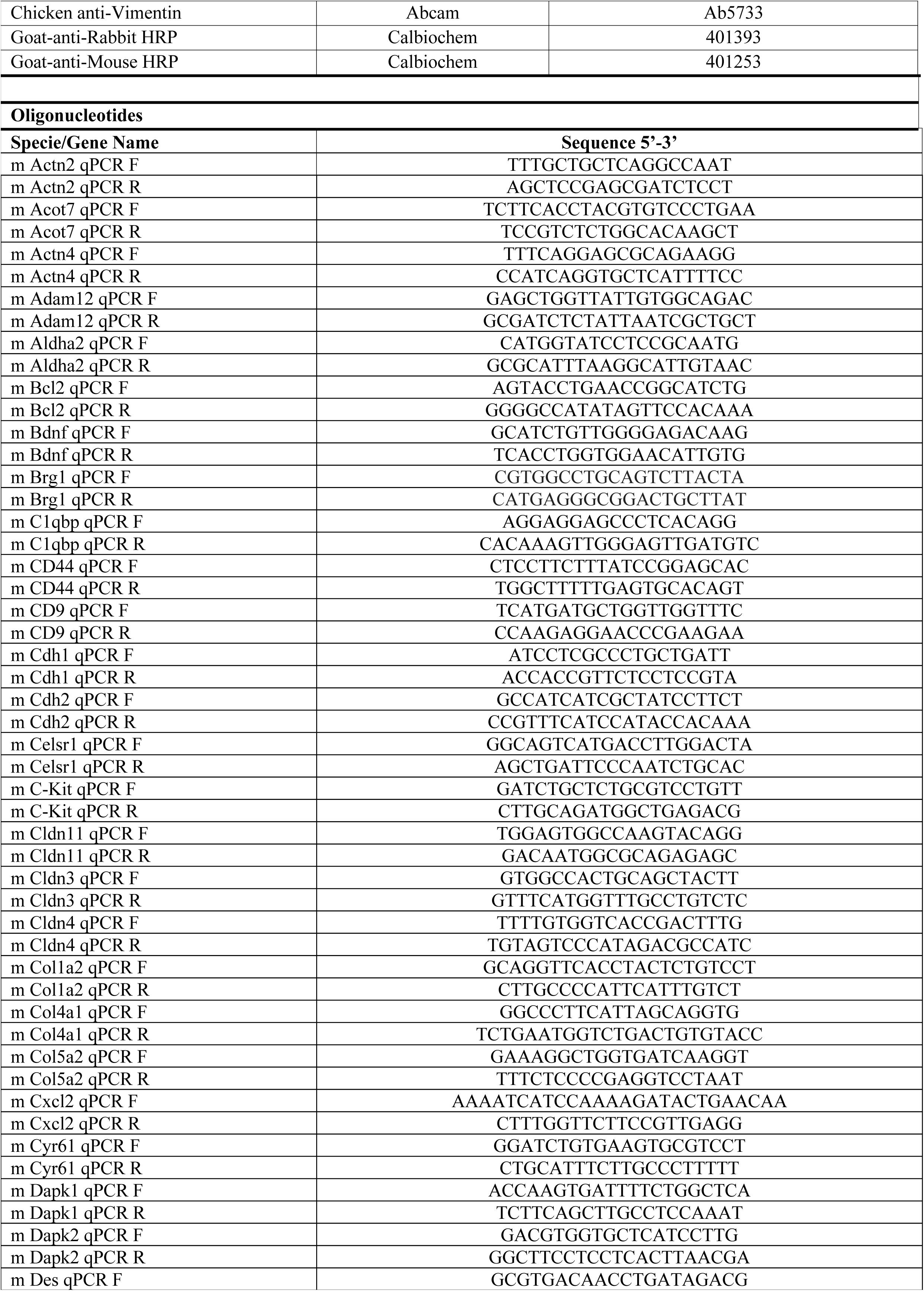
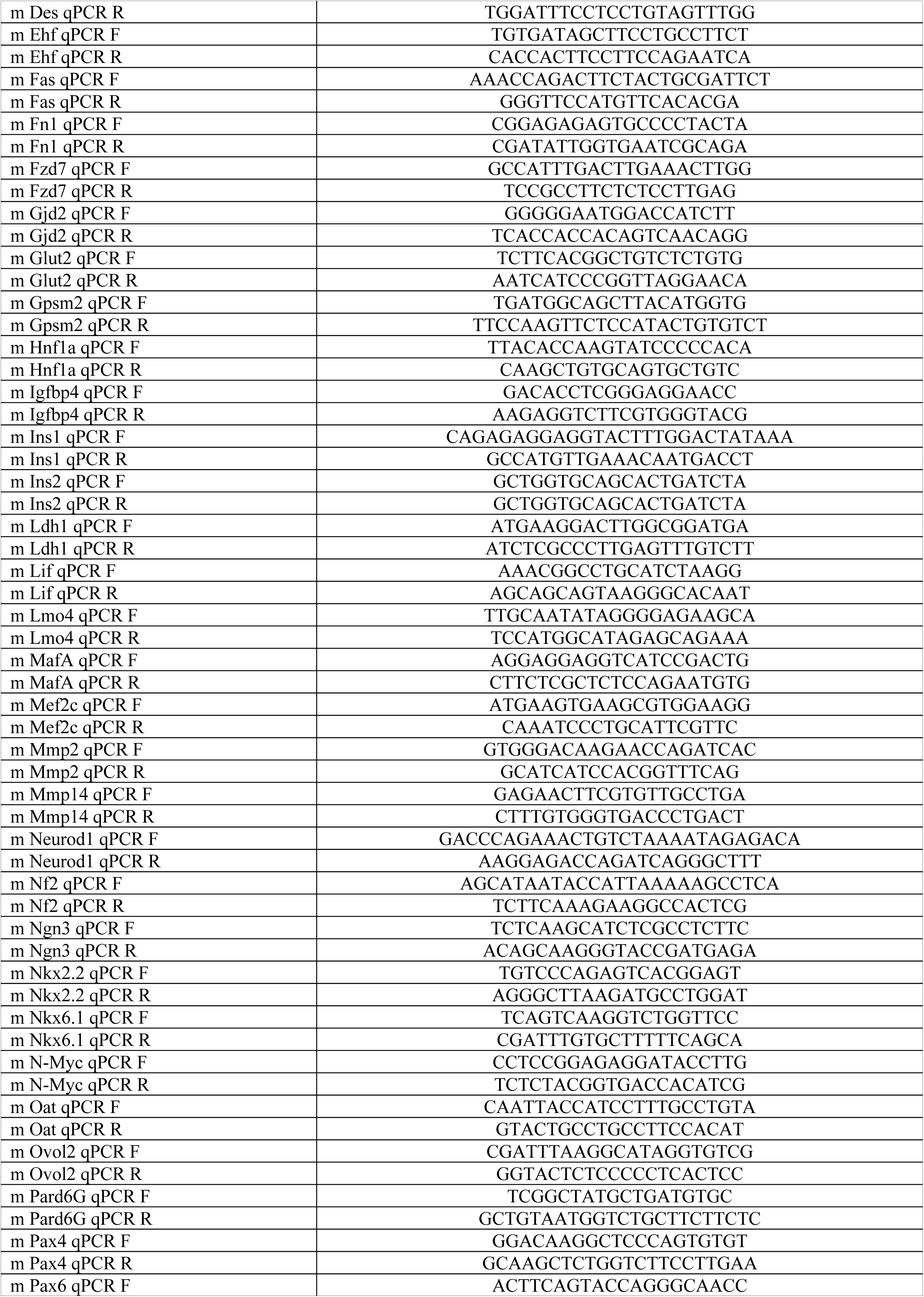
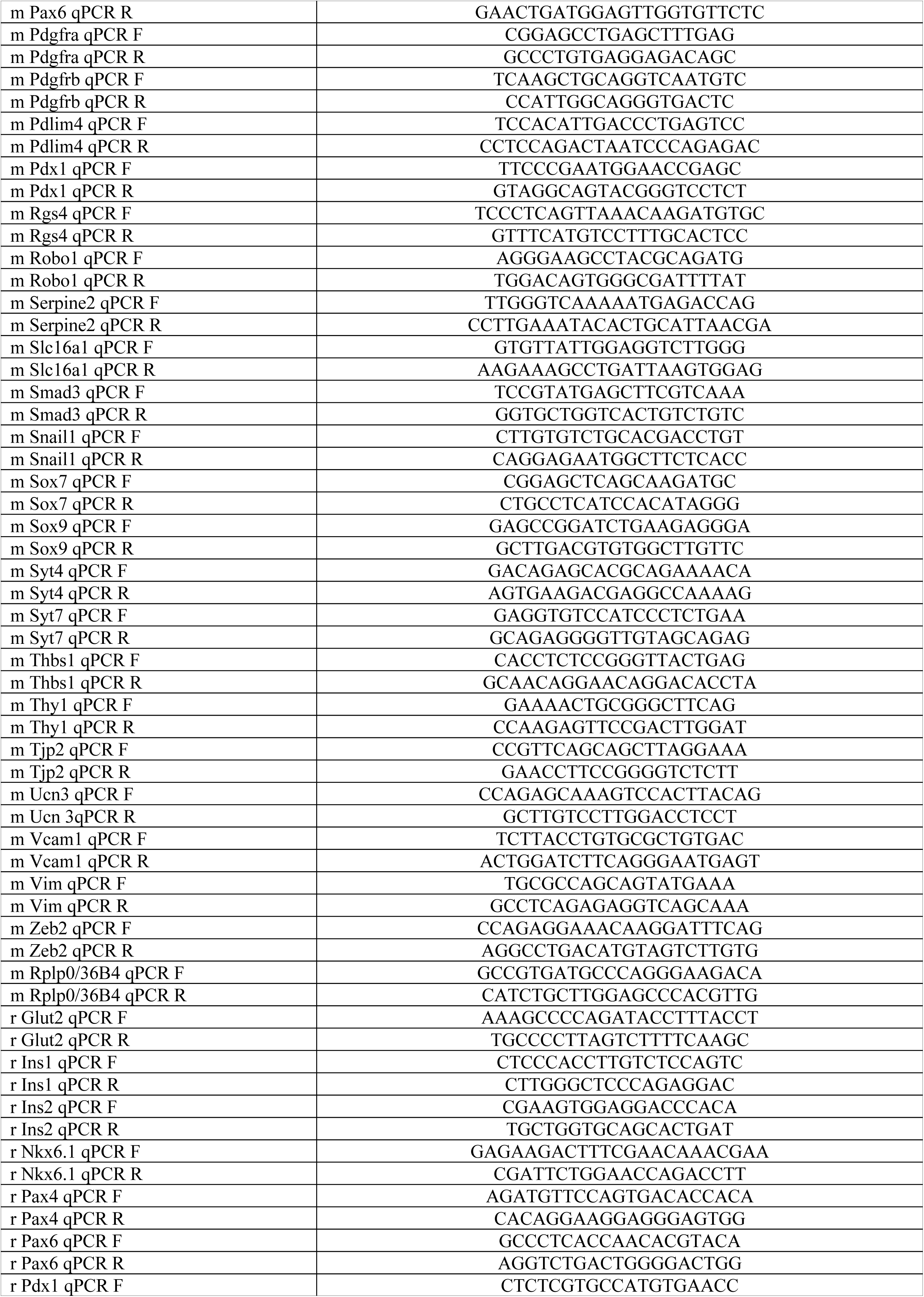
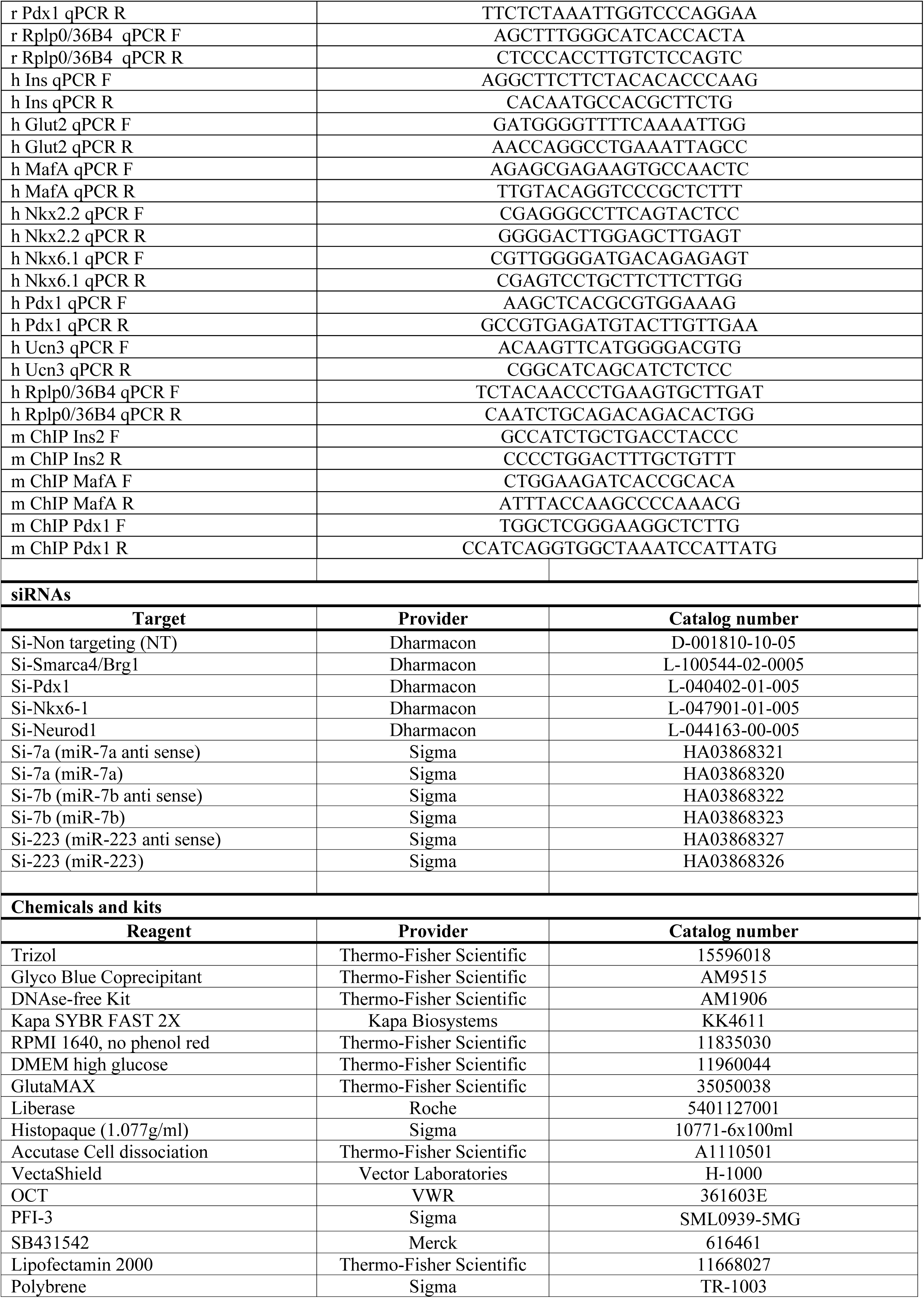
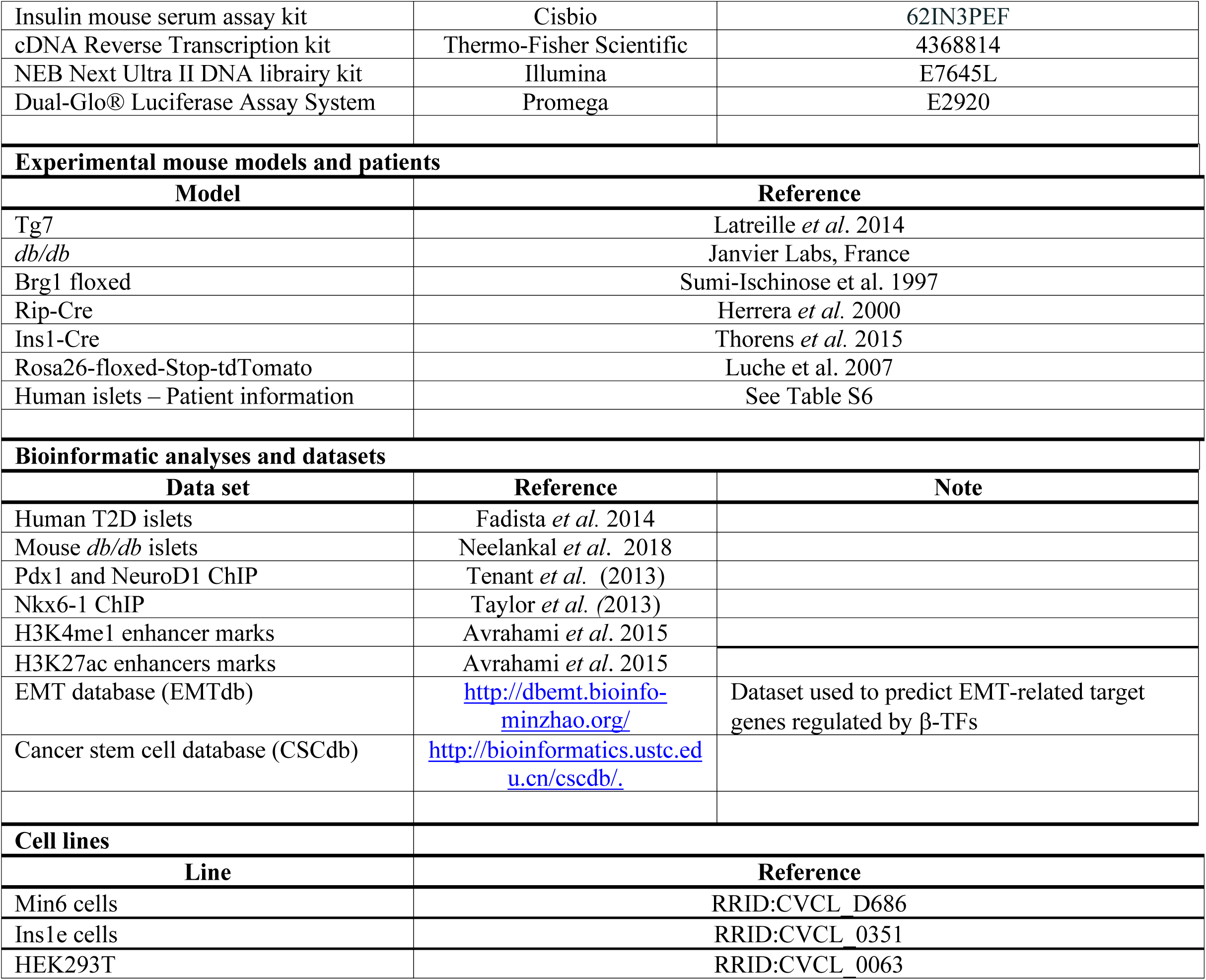

---

**Supplementary Figure 1.**
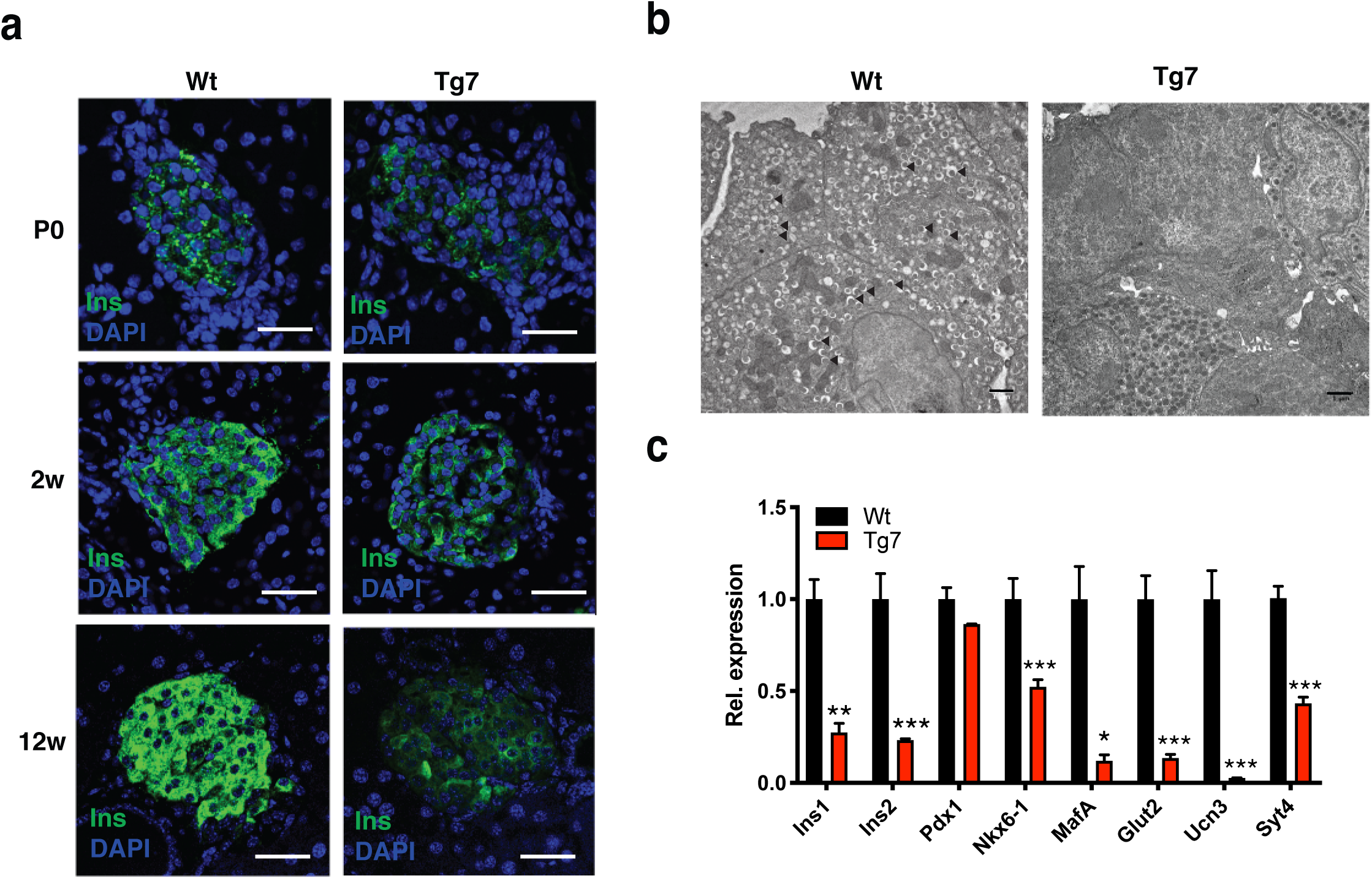
β-cell dedifferentiation in Tg7 mice. (A) Insulin immunofluorescence of pancreatic sections from wildtype (Wt) and Tg7 mice at birth (P0), 2-week (2w) and 12-week (12w) of age. Scale bar: 30μm (B) Electron microscopy of 12w Wt and Tg7 islet preparations. Arrowheads indicate the presence of insulin granules Scale bar: 1μm (C) Relative expression of β-cell identity genes in islets isolated from Wt and Tg7 mice at 12 weeks of age (n=6)

**Supplementary Figure 2.**
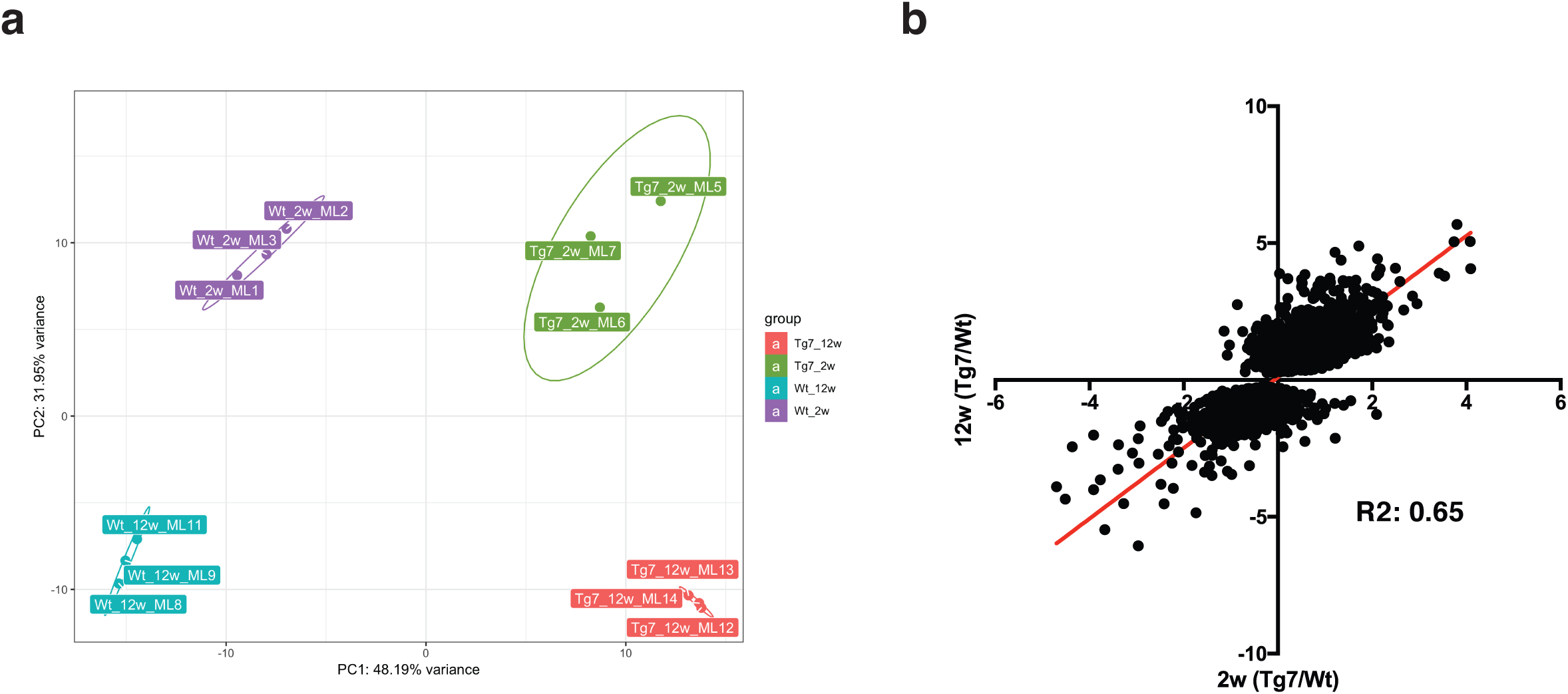
RNA-seq analysis in islets isolated from Tg7 and Wt controls mice. (A) Principal component plot (PCA) plot of the RNA-seq data obtained in islets isolated from 2 and 12 week old Wt (Wt-2w and Wt-12w) and Tg7 (Tg7-2w and Tg7-12w) mice. Each dot represents a sample. (B) Scatter plot of log2-transformed gene expression fold change in 2w versus 12w (Tg7/Wt) ratios. Each dot represents a gene

**Supplementary Figure 3.**
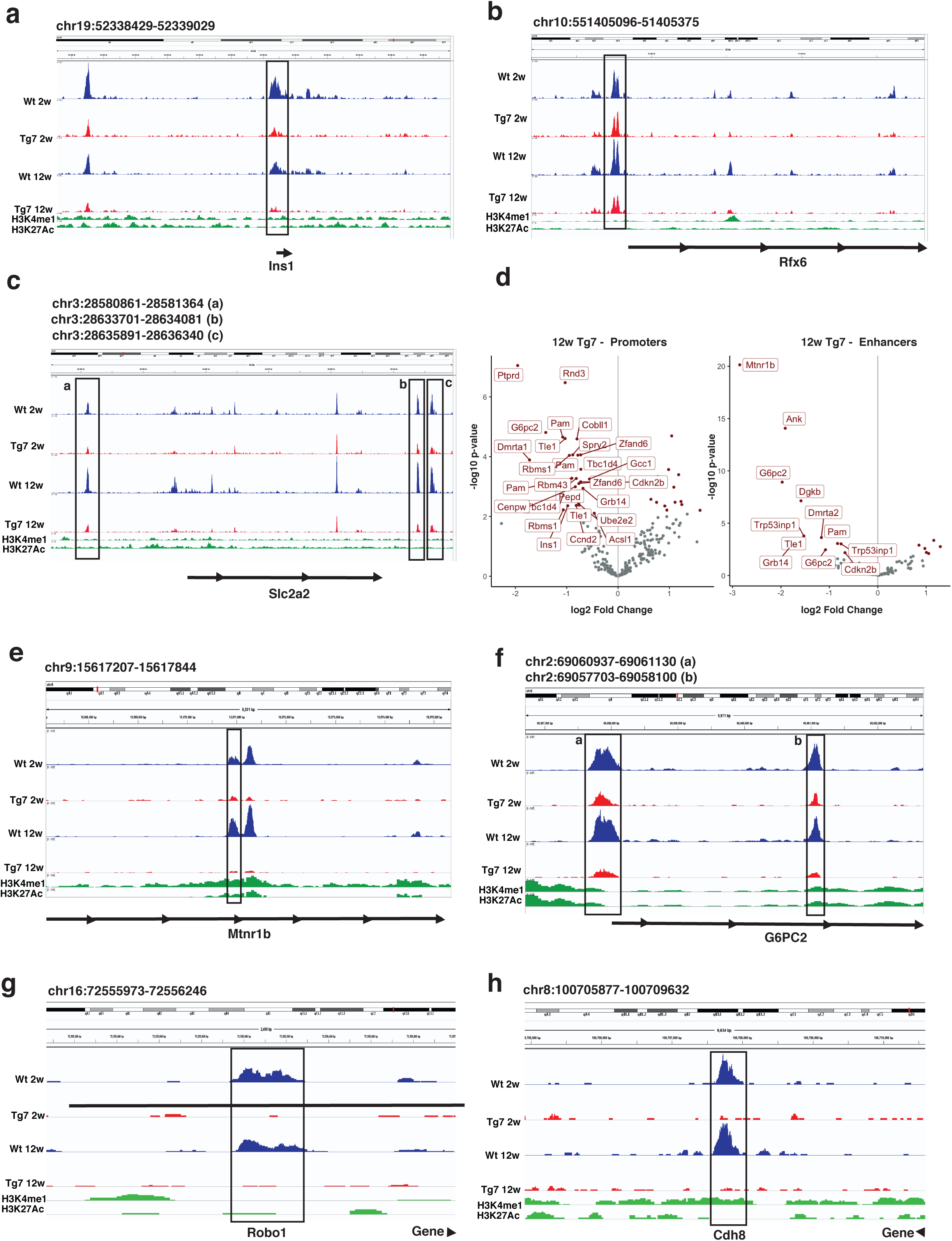
Closure of chromatin landscapes at β-cell-and epithelial-specific loci in dedifferentiated β-cells. (A-C) Representative screen shots depicting closing DAR associated to b-cell identity genes obtained in the ATAC-seq in islets from isolated from 2-and 12-week-old Wt (Wt-2w and Wt-12w) and Tg7 (Tg7-2w and Tg7-12w) mice. A) *INS1* locus; B) *RFX6* locus and C) *SLC2A2* locus. Data processed in IGV 2.4.11. pAdj<0.05 (D) Volcano plot of DARs in diabetes genome-wide association studies (GWAS) genes in 12w Tg7 mice and controls. Left: DAR found in promoters and Right: DARs found in H3K4me1^+^/H3K27Ac^+^ enhancers. Each dot represents a DAR identified in the ATAC-seq. DARs with differential accessibility in Tg7 versus Wt are depicted and labeled in red. (E-H) Representative screen shots depicting closing DAR associated to GWAS genes (E-F) and epithelial cell identity (G-H) identified in the ATAC-seq in islets isolated from Tg7 mice. pAdj<0.05

**Supplementary Figure 4.**
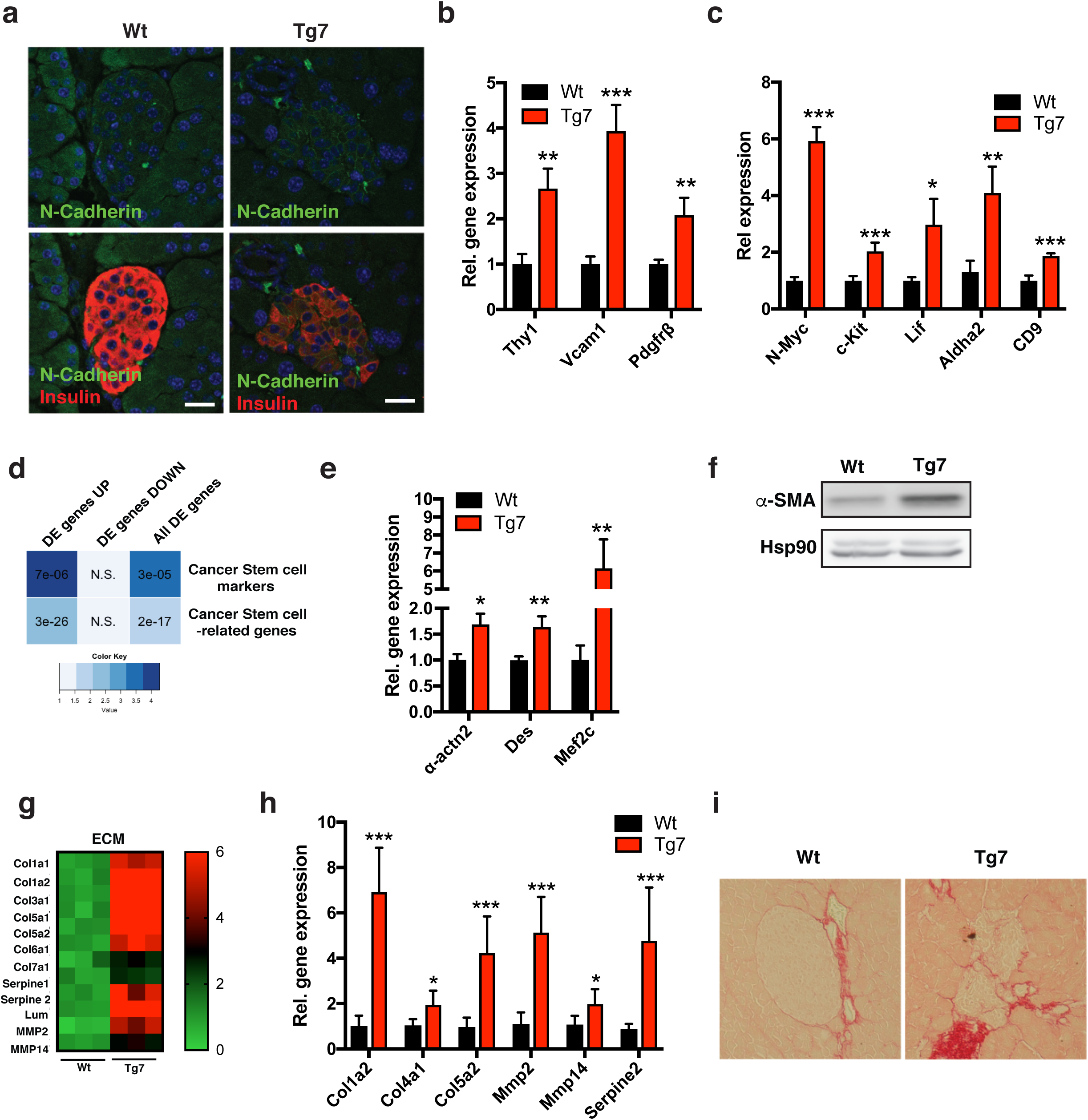
Induction of an EMT process in islets of Tg7 mice. (A) Cdh2/N-Cadherin (green) and insulin (red) co-immunofluorescence on pancreatic sections from 12-(12w) week-old Tg7 and Wt control mice. Scale bar: 30μm (B-C) mRNA levels of mesenchymal (B) and stemness (C) markers in islets from 12w Tg7 mice and controls (n=6). (D) Fisher exact test from LRT RNA-seq data demonstrating enrichment of differentially expressed genes in Tg7 islets (pAdj<0.01) with the Cancer Stem Cell and Cancer Stem cell-related gene sets; enrichment p-value corrected by Benjamin-Hochberg method (E) mRNA levels for myofibroblastic markers in islets from 12w Tg7 mice and controls (n=6) (F) Western blotting for α-SMA in 12w Tg7 islets normalized for the Hsp90 loading control. (G) Heat map of normalized DESeq2 counts in for indicated ECM components in Tg7 islets versus Wt controls; pAdj <0.05. (H) mRNA levels for indicated ECM components in islets from12w Tg7 mice and controls (n=6)(I) Sirius Red staining of pancreatic sections from from 12w Tg7 mice and controls. Unpaired Student’s t-test unless stated otherwise. Data are means ± SEM, *p < 0.05, ***p < 0.001

**Supplementary Figure 5.**
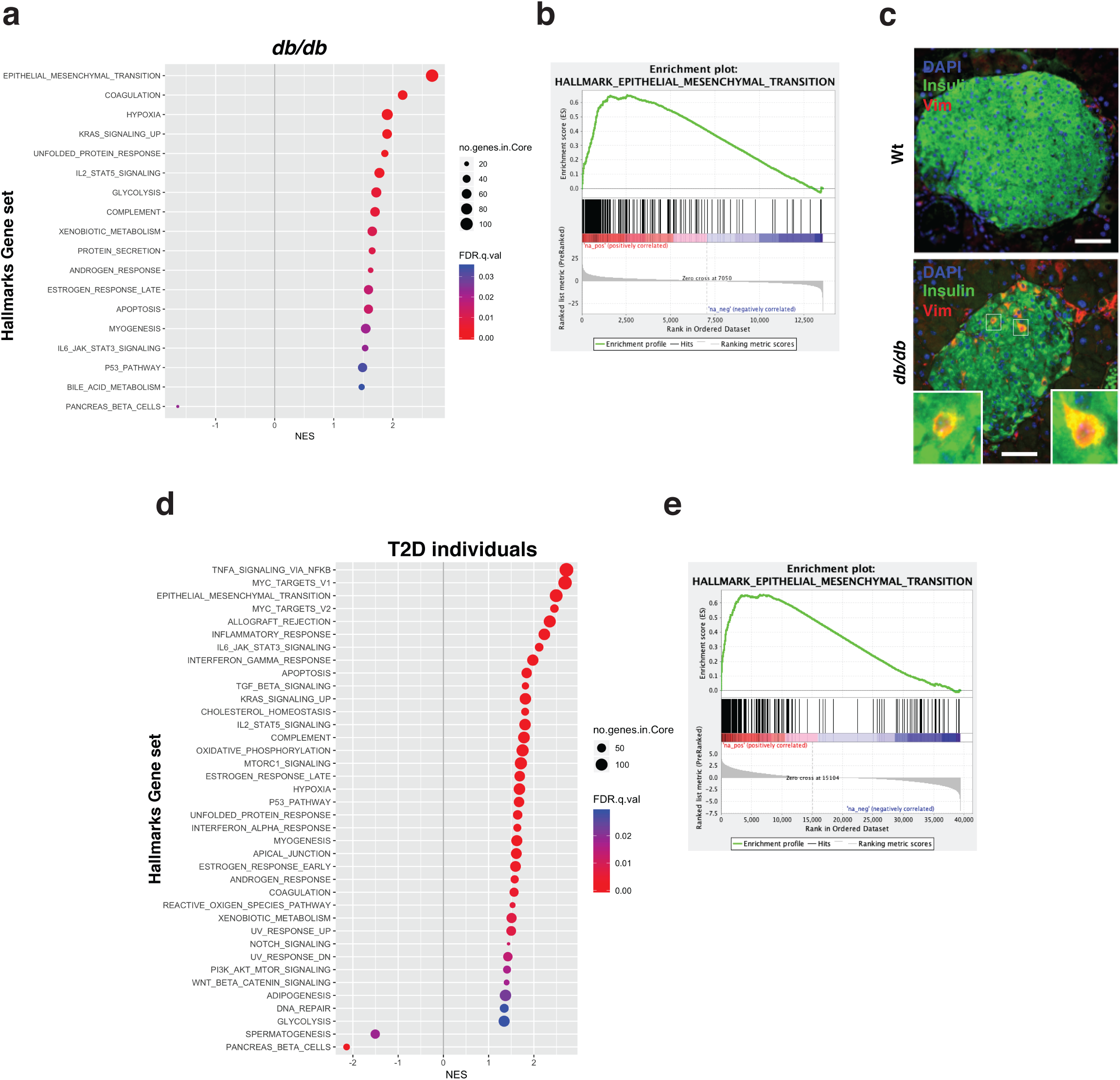
EMT genetic signature in islets from *db/db* mice and T2D individuals. Gene set enrichment analysis (GSEA) of MSigDB Hallmark genes in upregulated pre-ranked gene ratios (Tg:Wt) of islets isolated from *db/db* mice from Neelankal *et al.* (2018) (A-B) and T2D individuals from Fadista *et al.* (2014) C) Insulin (green) and Vimentin (Vim-red) immunofluorescence in pancreatic sections from 14 week old db/db mice. Nuclei (blue) revealed by DAPI staining. Scale bar: 40μm. (D-E). B and D panel are GSEA enrichment plots for the “epithelial to mesenchymal ransition” MSigDB hallmark gene in *db/db* mice and T2D individuals, respectively.

**Supplementary Figure 6.**
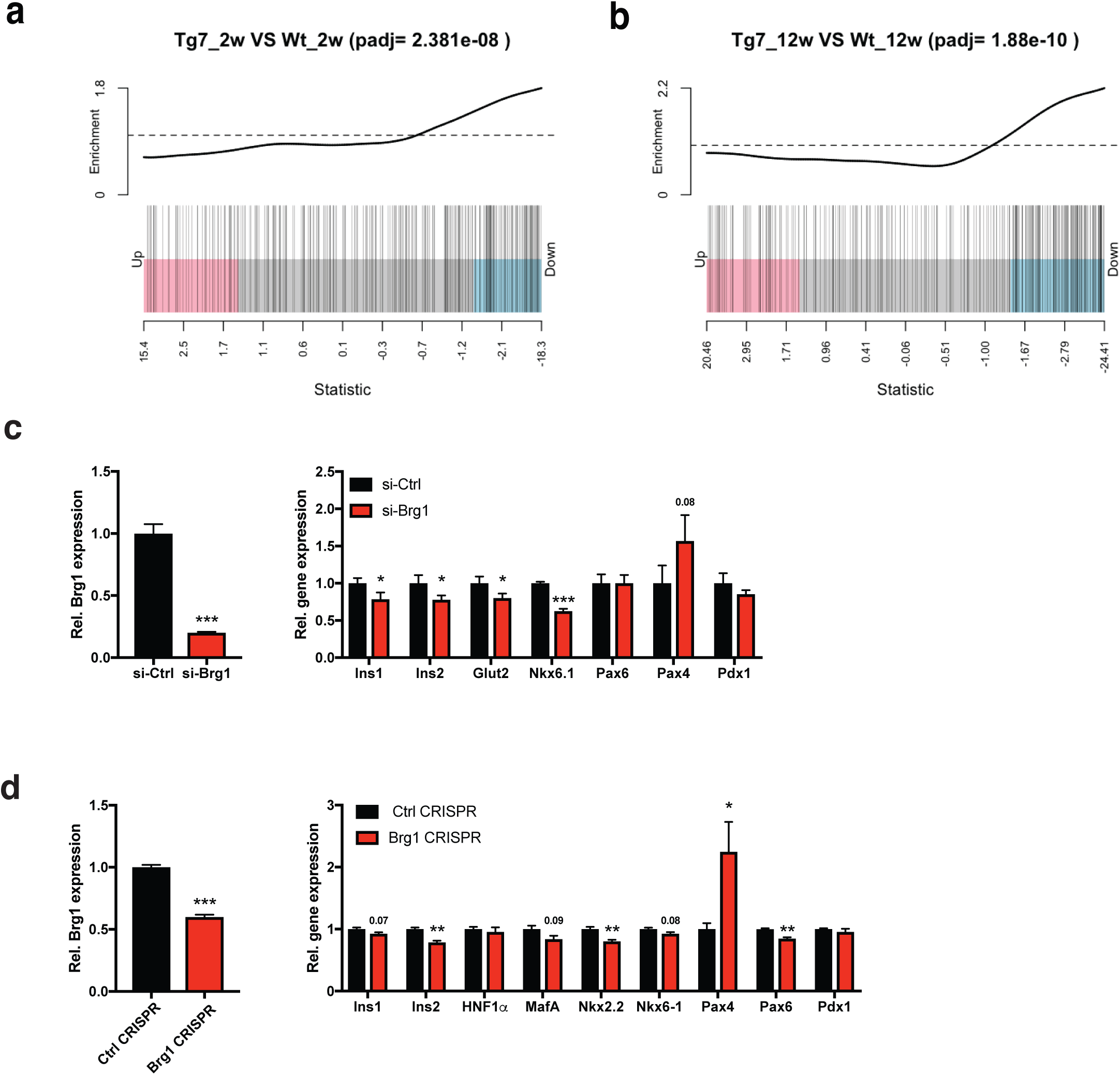
Smarca4/Brg1 is a target of miR-7 and is required to maintain β-cell identity. (A-B) Enrichment plot for miR-7 targets in islets from 2-(2w) (A) and 12-(12w) (B) Tg7 mice. List of predicted miR-7 targets were obtained from TargetScan (n=302) and processed using Limma/WilcoxGST package to determine whether these targets tend to be enriched in differentially downregulated or upregulated genes as compared to all other detected genes in Tg7 islets. (D) mRNA levels of Brg1 (left) and indicated b-cell-specific (right) genes in Ins1e cells transfected with siRNA against Brg1 (n=3) (D) mRNA levels of Brg1 (left) and indicated b-cell specific (right) genes in stable pools of MIN6 cells with deletion of the Brg1 gene following CRISPR/Cas9-mediated genome editing. (n=3) Unpaired Student’s t-test unless stated otherwise. Data are means ± SEM, *p <0.05,**p<0.01, ***p<0.001

